# ZNF121 recruits YTHDF2 to modulate mRNA stability

**DOI:** 10.64898/2026.02.18.706452

**Authors:** Giovanni L. Burke, Syed Nabeel-Shah, Shuye Pu, Nujhat Ahmed, Shahir M. Morcos, James D. Burns, Ahmed Abosen, Guoqing Zhong, Eric I. Campos, Jack F. Greenblatt

**Affiliations:** Department of Molecular Genetics, University of Toronto, Toronto, Canada; Terrence Donnelly Centre for Cellular and Biomolecular Research, University of Toronto, Toronto, Canada; The Peter Gilgan Centre for Research and Learning, The Hospital for Sick Children, Toronto, Canada

**Keywords:** m6A, YTHDF2, C2H2-Zinc Finger, ZNF121, RNA-binding, mRNA stability

## Abstract

*N*^6^-methyladenosine (m6A) is the most abundant internal modification in mRNA and has been shown to regulate gene expression through the binding of specific reader proteins, such as YTHDF2, which promotes mRNA decay. Previous studies indicate that YTHDF2 has relatively weak intrinsic RNA-binding affinity, suggesting that additional factors may facilitate its association with target transcripts. Here, we show that the C2H2-Zinc Finger protein, ZNF121, binds mRNA in cells and physically interacts with YTHDF2 in the cytoplasm. We demonstrate that approximately 80% of ZNF121-bound mRNAs are also YTHDF2 targets, and that their binding sites highly correlate with each other. Loss of ZNF121 impairs YTHDF2 binding to shared targets and increases their stability, independent of the presence of m6A modifications. Moreover, co-regulated transcripts are enriched for cell-cycle–related pathways, and ZNF121 depletion leads to elevated expression of the oncogene MDM2, implicating ZNF121 in growth control and the DNA damage response. Our findings identify ZNF121 as a cofactor that enhances YTHDF2-mediated mRNA regulation and reveal a previously unknown layer of control in mRNA decay.

## Introduction

Of the over 100 different RNA modifications that have been discovered (Boccaletto et al. 2018), *N^6^*-methyladenosine (m6A) is the most abundant and well-characterized mRNA modification. Functionally, m6A regulates co- and post-transcriptional mRNA processing, structure, stability, and translation efficiency, with biological consequences related to human development and several diseases, including cancer (Flamand et al. 2024; Roundtree et al. 2017).

m6A is deposited mostly sub-stoichiometrically at the DRACH consensus motif (D=A/G/U, R=A/G, H=A/C/U) and is enriched in 3’UTRs, with a peak around the stop-codon (Dominissini et al. 2012; Meyer et al. 2012). The METTL3-METTL14 methyltransferase complex (MTC) catalyzes m6A deposition via the methyltransferase activity of METTL3 (Liu et al. 2014). Additional components of the m6A-METTL-associated complex modulate MTC localization (Flamand et al. 2024). Conversely, FTO and ALKBH5 demethylate m6A, highlighting the dynamic reversibility and regulation of m6A (Jia et al. 2011; Zheng et al. 2013).

The cytosolic paralogues YTHDF1, YTHDF2, and YTHDF3 bind to m6A through their highly conserved YTH domains, which contain a tryptophan cage that selectively recognizes the m6A modification (Zhu et al. 2014; Xu et al. 2015, 2014; Li et al. 2014, 2020). The binding of these and other reader proteins modulates specific molecular processes, including translation, splicing and mRNA decay (Wang et al. 2014a; Du et al. 2016; Wang et al. 2015; Xiao et al. 2016), resulting in various effects on development (Collignon et al. 2023; Wang et al. 2014b). YTHDF2 specifically facilitates the decay of mRNA targets by recruiting the CNOT1 subunit of the CCR4-NOT poly(A)-tail decay complex (Wang et al. 2014a; Du et al. 2016).

Although YTH domains specifically bind m6A-modified mRNA, their binding is relatively weak. For YTHDF2, the K_d_ *in vitro* for m6A has been estimated at ∼2.5 µM (Zhu et al. 2014; Luo and Tong 2014; Theler et al. 2014; Xu et al. 2015, 2014; Meyer and Jaffrey 2017; Li et al. 2014). It has been estimated that approximately 25% of cellular mRNAs are methylated, and that these methylated transcripts on average harbour 1-3 m6A modifications each (Dominissini et al. 2012; Meyer et al. 2012; Liu et al. 2022; Perry et al. 1975; Desrosiers et al. 1974). Given ∼300,000 mRNA molecules per cell (with mean transcript length ∼2 kb), and an average of 1-3 m6A-modified sites per transcript (Dominissini et al. 2012; Meyer et al. 2012), the intracellular mRNA concentration of m6A is estimated to be ∼100-400 nanomolar. Considering the relatively weak intrinsic RNA-binding affinity of YTHDF2 and the widespread distribution of m6A sites across the transcriptome, YTHDF2-RNA interactions in cells may be dynamic and potentially influenced by additional factors. We therefore hypothesized that protein–protein interactions could enhance or stabilize YTHDF2 association with its target transcripts. Such cofactors could also help confer transcript selectivity among the highly similar YTHDF paralogues (Zaccara and Jaffrey 2020).

Cys2-His2 zinc finger proteins (C2H2-ZFPs), the largest class of DNA-binding transcription factors, with >700 proteins in humans, are characterized by their C2H2-zinc finger domains (ZnFs) (Klug 2010). While most C2H2-ZFPs are poorly characterized, they are generally diverse in their protein-protein interactions (PPIs) and DNA-binding preferences (Schmitges et al. 2016). They have also recently emerged as a large class of putative RNA-binding proteins (RBPs). Including classic examples like TFIIIA and YY1 (Seifart et al. 1989; Sigova et al. 2015), over 100 C2H2-ZFPs have been shown to directly bind RNA in cells through CLIP (UV crosslinking and immunoprecipitation)-based approaches and other methods in two independent studies (Nabeel-Shah et al. 2024b, Gosztyla et al. 2024).

C2H2-ZFPs appear to function in diverse RNA regulatory processes, ranging from alternative splicing to alternative cleavage and polyadenylation and mRNA stability (Han et al. 2017; Song et al. 2022). Some C2H2-ZFPs may be involved in gene regulation mediated by m6A modifications in mRNA. For example, we recently found that ZBTB48 is a C2H2-ZFP that interacts with the m6A eraser protein, FTO, facilitating the recruitment of FTO to methylated targets and modulating their expression (Nabeel-Shah et al. 2024a). Further, many C2H2-ZFPs, including ZNF784, which stabilizes some of its m6A-mRNA targets, bind mRNA around sites of m6A enrichment (Nabeel-Shah et al. 2024b).

ZNF121, a relatively poorly characterized C2H2-ZFP, stimulated our interest due to its potential role in cell proliferation and cancer progression via its characterized interactions with BRCA1 and MYC (Luo et al. 2018, 2016). ZNF121 contains eleven ZnFs and an *N-*terminal intrinsically disordered region (IDR), and we have previously shown that ZNF121 interacts with proteins involved in DNA repair, RNA localization, and cell cycle regulation (Schmitges et al. 2016; Nabeel-Shah et al. 2024b). Additionally, ZNF121 positively regulates *MYC* transcript abundance, and depletion of ZNF121 results in increased cell cycle arrest (Luo et al. 2018, 2016). Luo et al. (2018) showed that ZNF121 putatively interacts with components of the PAN2-PAN3 poly(A) decay complex, implicating it in RNA processing and metabolism. Given our recent findings that C2H2-ZFPs function in post-transcriptional regulation, and that ZNF121 interacts with certain RNA-binding proteins (e.g., SON, CARM1, AKAPL8, and PAN2) (Schmitges et al. 2016; Nabeel-Shah et al. 2024b; Luo et al. 2018), we aimed to further characterize the role of ZNF121 in mRNA regulation.

In this study, we examined how ZNF121 might act as an interacting cofactor for YTHDF2, finding that ZNF121 binds mRNA near YTHDF2 on common mRNA targets, including many related to the regulation of cell cycle. It also regulates the binding of YTHDF2 to these targets and target stability, independent of m6A.

## Results

### C2H2-ZFP ZNF121 interacts with YTHDF2

To begin characterizing ZNF121’s functions, we initially analyzed our previously published raw AP-MS data (Schmitges et al. 2016). To identify the full complement of ZNF121’s PPI partners, we filtered ZNF121’s AP-MS data against 25 GFP control purifications using Significance Analysis of Interactome (SAINTexpress) (Teo et al. 2014). Application of SAINTexpress identified several interaction partners that passed our statistical threshold of Baysian false discover rate (BFDR) ≤ 0.01 (Supplemental Table 1). Consistent with ZNF121’s previously known interactions with RBPs, we identified significant interaction partners enriched in RNA-related pathways, including RNA processing, mRNA metabolism, translation, and splicing (Supplemental Fig S1A). Interestingly, SAINTexpress analysis also identified the m6A reader protein, YTHDF2, as one of the significant interaction partners of ZNF121 (BFDR ≤ 0.01), suggesting that ZNF121 might have a role in m6A metabolism. We, therefore, focused our attention to further characterizing the functional relevance of the ZNF121-YTHDF2 interaction.

To validate ZNF121’s interaction with YTHDF2, immunoprecipitation (IP) with anti-GFP antibodies was performed in HEK293 cells expressing *N-*terminally GFP-tagged ZNF121. Compared to control IPs in cells expressing GFP alone, endogenous YTHDF2 co-purified specifically with GFP-ZNF121 (Supplemental Fig. S1B). This interaction was further substantiated by reciprocal co-IP experiments (Supplemental Fig. S1C). To exclude the possibility that the observed interaction was driven by overexpression of GFP-ZNF121, we performed IP of endogenous YTHDF2 from whole-cell lysates of wild-type HEK293, HeLa, MCF-7, and HCT-116 cells, and probed for endogenous ZNF121. These cell lines were selected to represent distinct tissue origins and genetic backgrounds, as well as ZNF121’s previously observed role in cancer (Luo et al. 2018, 2016). In all cases, ZNF121 co-precipitated with YTHDF2, demonstrating that this interaction occurs at endogenous levels across multiple cell types (Fig. 1A). Of note, all samples were treated with Benzonase nuclease prior to IPs, making it unlikely that these interactions were mediated by DNA or RNA. We also subjected lysates from cells expressing GFP-ZNF121 to increasing amounts of RNase I, followed by co-IP, where we observed little change in ZNF121’s ability to co-precipitate YTHDF2 (Supplemental Fig. S1D).

**Figure 1.**
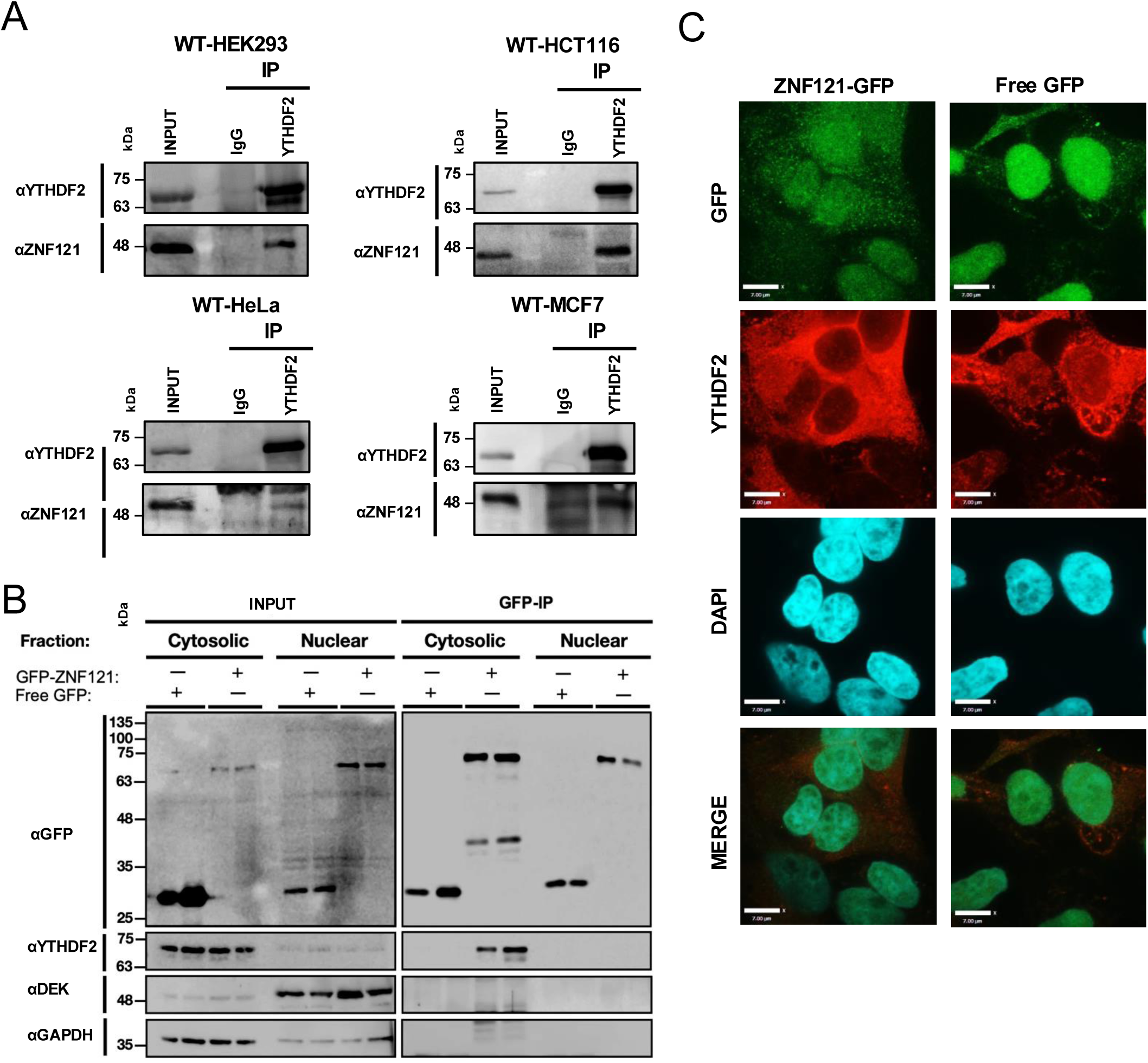
ZNF121 and YTHDF2 interact in the cytoplasm. (*A*) Western blots showing co-IP of endogenous ZNF121 after IP of endogenous YTHDF2 with anti-YTHDF2 antibodies from extracts of the indicated cell types. IgG was used as a negative control antibody. (*B*) Western blots showing co-IP of YTHDF2 after IP with anti-GFP antibodies from nuclear and cytoplasmic cell fractions of HEK293 cells expressing either GFP or GFP-ZNF121. DEK and GAPDH were used as nuclear and cytoplasmic markers, respectively. (*C*) Immunofluorescence analysis utilizing anti-GFP and anti-YTHDF2 antibodies in HEK293 cells expressing GFP or GFP-ZNF121. Scale bar = 7 μm at 100x magnification.

To characterize the localization of ZNF121 relative to cytoplasmic YTHDF2, we performed subcellular fractionation experiments using extracts from cells expressing GFP-ZNF121. GFP-ZNF121 was found in both the nucleus and cytoplasm, whereas YTHDF2 was present only in the cytoplasm, as reported previously by (Wang et al. 2014a) (Supplemental Fig. S1E). DEK, a protein involved in chromatin organization (Hu et al. 2007), and GAPDH were used as nuclear and cytosolic marker proteins, respectively. To ensure that GFP-ZNF121 was not driven to the cytoplasm solely by overexpression, this experiment was also carried out with WT-HEK293 cells using anti-ZNF121 antibodies, producing similar observations (Supplemental Fig. S1F). Utilizing nuclear and cytosolic fractions from HEK293 cells expressing either free GFP or GFP-ZNF121, IPs with anti-GFP were also performed, confirming that YTHDF2 only interacts with cytoplasmic GFP-ZNF121 (Fig. 1B). As well, immunofluorescence microscopy confirmed that GFP-ZNF121 is diffuse throughout the cytoplasm and nucleus (Fig. 1C).

### Identifying the ZNF121-YTHDF2 protein interaction surfaces

While the structure of the YTH domain of YTHDF2 has been experimentally determined (Zhu et al. 2014; Li et al. 2014), the structure of ZNF121 has not yet been characterized. We therefore predicted the structure of ZNF121 using AlphaFold 2.0 (Jumper et al. 2021). The resulting model indicates that the globular ZnF portion adopts a right-handed helical configuration, with high Predicted Local Distance Difference Test (pLDDT) and low predicted alignment error (PAE) across individual zinc finger motifs (Supplemental Fig. S2A-C). Subsequently, we used AlphaFold-Multimer (Evans et al. 2022) to predict the co-structure of ZNF121 and YTHDF2 (Fig. 2A, Supplemental Fig. S2A-D). As a measure of the reliability of the structures generated by AlphaFold 2.0, we aligned the predicted YTHDF2 structure with its structure determined by X-ray diffraction (Li et al., 2014) (Supplemental Fig. S2E), finding that the two structures align very well. Interestingly, the m6A binding pocket of YTHDF2 remains unobstructed in the co-structure, suggesting YTHDF2 could still potentially interact with RNA and m6A when in complex with ZNF121.

**Figure 2.**
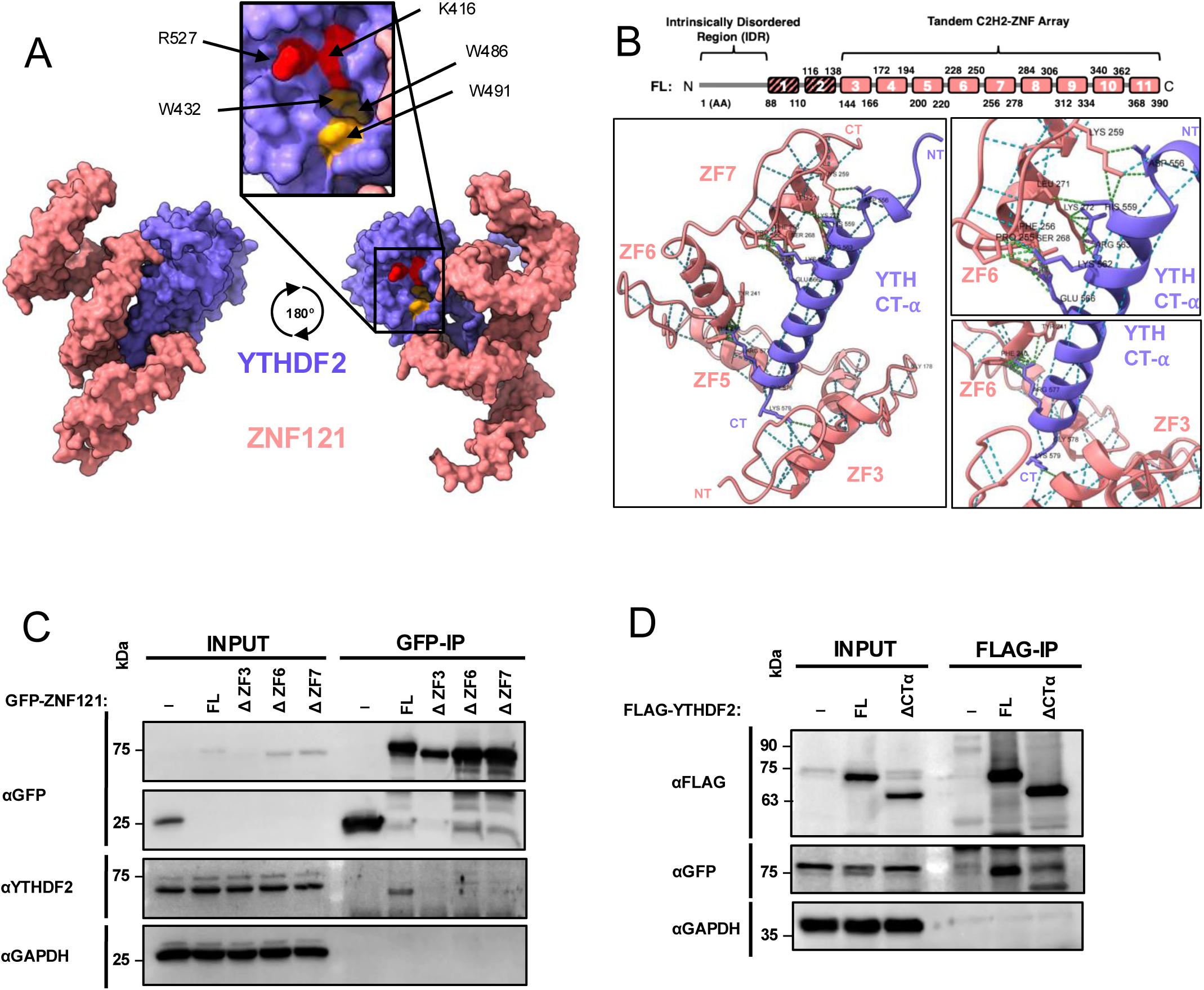
The C-terminal region of YTHDF2 facilitates an interaction with the ZnFs of ZNF121. (*A*) Space-filling model of the ZNF121-YTHDF2 co-structure predicted by AlphaFold multimer, using structures for ZNF121 (pink) and YTHDF2 (purple) predicted by AlphaFold 2.0. Only the interacting globular domains are depicted for clarity. Highlighted are the tryptophan cage residues of YTHDF2 that bind m6A (yellow) and that contact the RNA backbone (red). (*B*, *top*) Protein organization of ZNF121, where boxes indicate ZnFs. (*Bottom*) Simplified ribbon co-structure of ZNF121 and YTHDF2 on the left, highlighting the predicted points of contact (green dashes) between the C-terminal α-helix (CTα) of YTHDF2 and ZnFs 3, 6 and 7 of ZNF121. (*Right*) Two zoomed-in perspectives. Contacts are atom pairs displayed as pseudobonds with a Van Der Waals overlap of ≥ 0.4 Å between the two proteins. Hydrogen bonds, displayed as blue dashed lines, are displayed as pseudobonds with a distance tolerance of 0.4 Å between peptide chains and an angle tolerance of 20° (default settings). Purple dashed lines represent predicted clashes using default settings. Predictions were generated by ChimeraX (Meng et al., 2023). (*C*) Western blots showing co-IP of YTHDF2 after IP with anti-GFP antibodies from extracts of HEK293 cells expressing GFP-ZNF121 fusion proteins that are full length (FL) or lack ZNFs 3 (ΔZF3), 6 (ΔZF6), or 7 (ΔZF7). (*D*) Western blots showing co-IP of GFP-ZNF121 after IP using FLAG antibodies of FLAG-YTHDF2 that is full length (FL) or lacks the C-terminal residues 550-579 of YTHDF2 (ΔCTα).

The folded structure of residues T550-G579, corresponding to the final α-helix of YTHDF2 at its *C*-terminus (CTα), was predicted by AlphaFold 2.0 but not resolved in experimentally determined structures (Zhu et al. 2014; Li et al. 2014) (Supplemental Fig. S2E,F). Expression of various tagged fragments of ZNF121 or YTHDF2 in cells followed by co-IP revealed that the *N-*terminal (NT) region of FLAG-ZNF121, which contains an 88 residue IDR and ZnFs 1-3, is sufficient for weak binding to YTHDF2 (Supplemental Fig. S2G,H). The IDR of ZNF121 is not, however, needed for its YTHDF2 interaction, since a co-IP performed with extracts from cells expressing full-length (FL) or IDR-truncated (ΔIDR) FLAG-ZNF121 revealed no change in the interaction with YTHDF2 (Supplemental Fig. S2I). Interestingly, the CTα is predicted to contact ZnFs 3, 6, and 7 of ZNF121 using the molecular visualization program, ChimeraX (Meng et al. 2023) (Fig. 2B). Consistently, deleting each of these ZnFs substantially reduces the interaction of GFP-ZNF121 with FLAG-YTHDF2 (Fig. 2C), suggesting that each of these ZnFs might be important for the interaction. Alternatively, deleting ZnF 6 or 7 may reposition downstream ZnFs so that they interfere with the interaction. These experiments suggested that the interaction of ZNF121 with YTHDF2 might require ZnF 3 and be stabilized by ZnFs 6 and 7.

Conversely, YTHDF2 residues 396-579 tagged with GFP, containing the YTH domain, is sufficient for binding to ZNF121 (Supplemental Fig. S2G,J). Moreover, deleting the CTα (ΔCTα) attenuates the interaction of FLAG-YTHDF2 with GFP-ZNF121 (Fig. 2D), suggesting that the CTα might stabilize the ZNF121-YTHDF2 interaction, as predicted by AlphaFold-Multimer analysis. Consistent with this observation, YTHDF1 and YTHDF3, whose proximal CTα sequences are virtually identical to that of YTHDF2 (Supplemental Fig. S2F), also bind ZNF121 in co-IP experiments (Supplemental Fig. S2K). Together, the data indicates that the CTα of the YTH domain of YTHDF2 and *N-*terminal ZnFs of ZNF121 are important for the ZNF121-YTHDF2 interaction.

### ZNF121 binds to mRNA and co-localizes on mRNA with YTHDF2

We next used the CLIP method to assess the RNA-binding capacity of ZNF121 in cells. Lysates from UV-treated cells expressing GFP-ZNF121 were subjected to varying degrees of RNA fragmentation with RNase I, followed by IP with anti-GFP, labeling with ^32^P, SDS-PAGE, and autoradiography. The protein-RNA complexes migrated more slowly than ZNF121 and were affected by the RNase I concentration, with reduced amounts of RNase I resulting in further reductions in migration due to the increased size of the protein-RNA complex (Fig. 3A).

**Figure 3.**
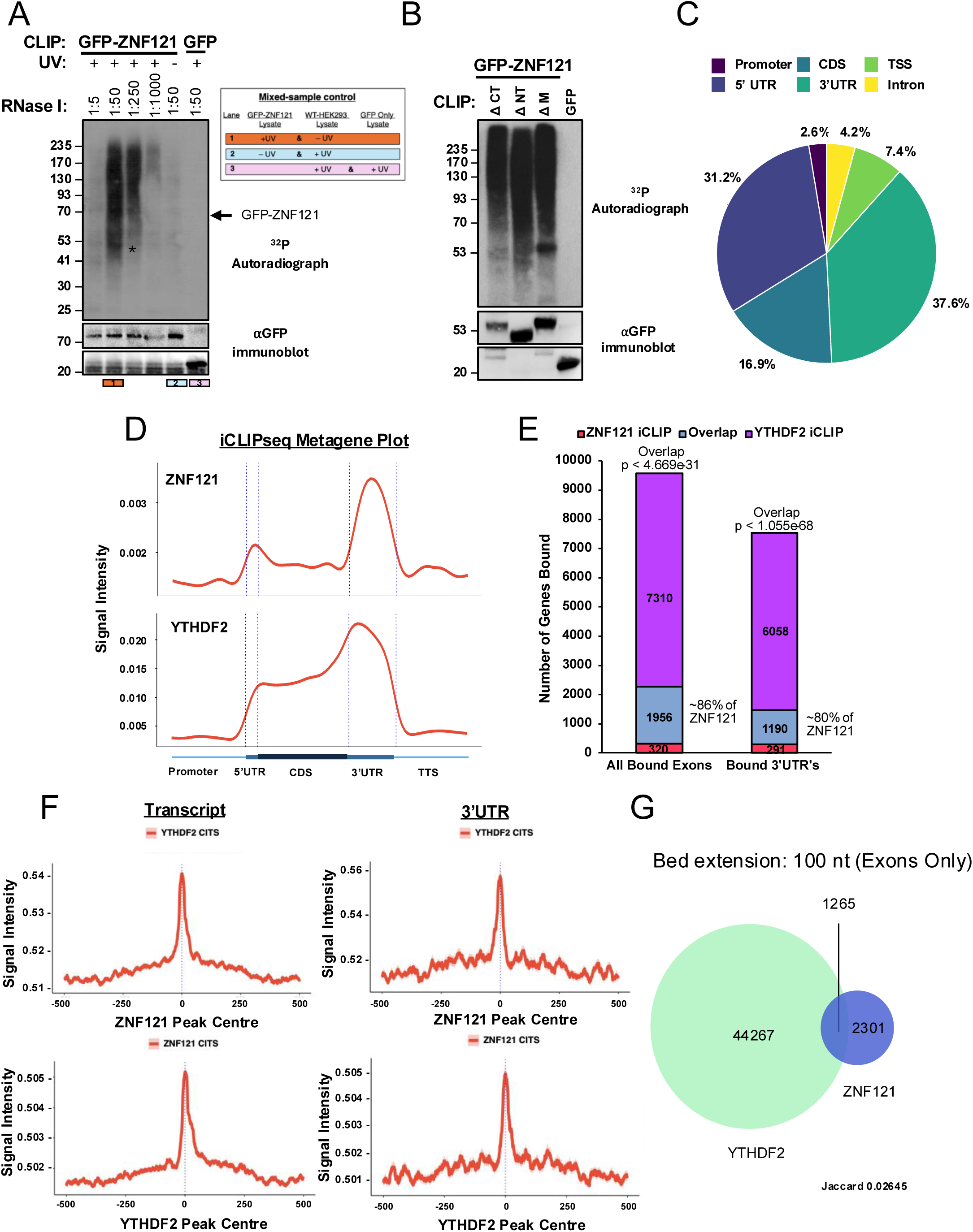
iCLIPseq identifies ZNF121 to primarily bind 3’UTRs near YTHDF2 binding sites. (*A*) CLIP-autoradiograph and Western blots derived from extracts of HEK293 cells expressing GFP or GFP-ZNF121, with or without UV treatment. For the CLIP autoradiograph, extracts were treated with the indicated concentrations of RNase I, followed by IP with anti-GFP, end-labeling of the RNA with ^32^P and SDS-PAGE. The asterisk (*) marks the position of IgG heavy chain and an arrow the position of ZNF121. Lanes 1, 2 and 3 (as indicated below in the Western blots and on the top right) indicate various experimental conditions: 1) lysate from UV-treated GFP-ZNF121-expressing cells mixed with lysate from non-UV-treated WT-HEK293 cells; 2) lysate from non-UV-treated GFP-ZNF121-expressing cells mixed with lysate from UV-treated WT-HEK293 cell lysate; and 3) lysate from non-UV-treated WT-HEK293 cells mixed with lysates from UV-treated GFP-expressing cells. (*B*) CLIP-autoradiogram and Western blot derived from extracts of UV-treated HEK293 cells expressing GFP or the indicated GFP-ZNF121 fusion proteins, after IP with anti-GFP, end-labeling of the RNA with ^32^P, and SDS-PAGE. ΔNT refers to a deletion of the IDR and ZnFs 1-3, ΔM refers to the deletion of ZnFs 4-7, and ΔCT refers to the deletion of ZnFs 8-9. Below are the corresponding Western blots probing with anti-GFP. (*C*) Pie chart depicting the percentage of GFP-ZNF121 CITS localizing to the indicated mRNA features after length normalization of the indicated features. (*D*, *Top*) Metagene plots of GFP-ZNF121 CITS, or (*Bottom*) YTHDF2 CITS, shown as signal intensity over input. (*E*) Overlap (blue) of transcripts bound by ZNF121 (red) and YTHDF2 (purple), as identified by iCLIPseq, either in all exons or only in their 3’UTRs. *p* determined by the hypergeometric test. (*F, Top*): Distribution of CITS derived from YTHDF2 iCLIPseq data around ZNF121 CITS in all exons (*left*) or only in the 3’UTR (*right*). (*Bottom*) Reciprocal analysis showing the distribution of CITS derived from ZNF121 around those of YTHDF2. (*G*) Overlap of YTHDF2 CITS and ZNF121 CITS located within 100 nt of each other in exons. Jaccard similarity coefficient was used to determine the degree of similarity between the two data sets.

To rule out the possibility that the observed RNA signal might be originating from *in vitro* ZNF121-RNA associations, we performed CLIP experiments using: 1) lysate from UV-treated cells expressing GFP-ZNF121 mixed with lysate from *non*-UV-treated WT-HEK293 cells; 2) lysate from *non-*UV-treated cells expressing GFP-ZNF121 mixed with lysate from UV-treated WT-HEK293 cells; and 3) lysate from UV-treated WT-HEK293 cells mixed with lysate from UV-treated cells expressing only GFP (all subjected to RNase I treatment) (Fig. 3A). Importantly, a radioactive RNA signal is observed only when lysate from UV-treated cells expressing GFP-ZNF121 is mixed with lysate from *non*-UV-treated WT-HEK293 cells. To rule out whether the presence of non-specific proteins co-purifying with ZNF121 in our CLIP experiments was a factor in the observed RNA signal, the precipitated proteins in CLIP and co-IP protocols were compared by silver staining after IP of GFP-ZNF121, revealing that CLIP produces much less protein background than standard co-IP conditions (Supplemental Fig. S3A). Together, these results indicated that co-precipitation of non-specific RBPs has minimal contribution to the observed RNA signal and that ZNF121 binds to RNA independent of other RBPs.

Since ZNF121 is a putative transcription factor, the observed radioactive signal could also be due to DNA. To this end, an over-digestion assay was performed, similar to our previous studies (Nabeel-Shah et al. 2024b). Briefly, UV-treated GFP-ZNF121-expressing HEK293 cells were lysed and processed according to the CLIP protocol. The IP’d protein-RNA complex was then subjected to an additional round of treatment with a high amount of RNase I or DNase. The RNA signal following SDS-PAGE and autoradiography was lost only upon additional RNase I, but not DNase, treatment, indicating that the radioactive CLIP signal is due to the crosslinking of ZNF121 to RNA, not DNA (Supplemental Fig. S3B). These results collectively suggested that the ZNF121’s DNA-binding has little contribution to the observed RNA signal in our CLIP experiments.

To identify the region of ZNF121 important for RNA-binding, three deletions to the full-length (FL) GFP-ZNF121 protein were made: 1) an *N-*terminal deletion (ΔN), 2) a “Middle” deletion (ΔM), and 3) a *C-*terminal (ΔCT) deletion (Supplemental Fig. S3C). These GFP-ZNF121 constructs were expressed from the HEK293 Flp-In™T-REx locus, followed by UV-treatment of the cells and CLIP-autoradiography. All three deletion mutants of GFP-ZNF121 crosslinked to RNA, and so neither the IDR nor any specific set of ZNFs is solely responsible for binding to RNA (Fig. 3B).

For a global, transcriptome-wide assessment of ZNF121’s mRNA binding, individual nucleotide resolution CLIP followed by sequencing (iCLIPseq) was performed on five biological replicates, utilizing size-matched inputs as a control. Only crosslink-induced truncation sites (CITS) that passed a threshold of FDR ≤ 0.01 were considered for downstream analysis (Supplemental Table 2). A total of 17,461 CITS peaks were identified, with ∼ 74% found in protein-coding transcripts (Supplemental Fig. S3D). ZNF121 predominantly crosslinked to mRNA in the 3’UTR and coding sequence (37.6% and 31.2% of CITS, respectively) after normalizing for feature length (Fig. 3C). Metagene analysis recapitulated ZNF121’s preference for 3’UTRs and showed that its distribution on mRNA is similar to that previously published for YTHDF2 (Nabeel-Shah et al. 2024b) (Fig. 3D).

To examine whether RNA-binding by ZNF121 correlates with its DNA-binding, we plotted ZNF121’s iCLIPseq CITS peaks around its previously reported ChIPseq peaks (Schmitges et al. 2016a) (as compared to random sites within 200 nt of the ZNF121 iCLIPseq CITS peak). We found there is no correlation between ZNF121’s iCLIPseq and ChIPseq peaks (Supplemental Fig. S3E). Moreover, when ZNF121’s iCLIPseq target genes and ChIPseq target gene promoters were compared, only ∼25% of ZNF121’s RNA-binding targets are also bound by ZNF121 on their corresponding promoter DNA (Supplemental Fig. S3F), and this overlap is not statistically significant (*p<*0.189). These analyses indicated that ZNF121 has distinct RNA- and DNA-binding preferences, with little overlap. Thus, the RNA-binding role of ZNF121 appears to be mostly or entirely decoupled from transcription.

We also examined whether ZNF121 prefers any particular mRNA features for binding. Utilizing the Discriminative Regular Expression Motif Elicitation (DREME) motif analysis suite (Bailey 2011) and iCLIPseq data, (Supplemental Fig. S3G), we found that the top predicted motifs are all U-rich, although their frequency of occurrence is relatively low (i.e., <10% for individual motifs). We thus also investigated whether ZNF121 might recognize specific mRNA structures. We utilized *in vivo* and *in vitro* generated icSHAPE and RNAplFold – an *in silico* method for RNA structure prediction (Flynn et al. 2016; Lorenz et al. 2011) – which have been used to investigate RNA-binding by C2H2-ZFPs and other well-characterized RBPs (Nabeel-Shah et al. 2024b). Scores with these methods indicate the probability of unpaired nucleotides at or surrounding the crosslinking sites of an RBP. The crosslinking sites of ZNF121 prefer, on average, unpaired RNA relative to the surrounding region of low unpaired probability based on *in vivo* and *in vitro* icSHAPE scores, as well as *in silico* RNA-plFOLD (Supplemental Fig. S3H). These results are similar to those for most C2H2-ZFPs analyzed in our previous study (Nabeel-Shah et al. 2024b) and indicate reaction with locally unpaired sequences within structured RNA.

Since ZNF121 binds to mRNA and interacts with YTHDF2, we determined to what extent ZNF121 targets are shared with YTHDF2. Specifically, we compared the iCLIPseq targets of ZNF121 and YTHDF2 and identified targets common to both proteins (YTHDF2∩ZNF121). Remarkably, 1,956 of the identified ZNF121 iCLIPseq mRNA targets (86%) are also YTHDF2 targets, with ∼80% specifically in 3’UTRs. On the other hand, only ∼ 22% of identified YTHDF2 targets (and ∼16 % specifically in 3’UTRs) are also bound by ZNF121 (Fig. 3E, Supplemental Table 3). This indicated that almost all ZNF121 targets are also YTHDF2 targets, whereas only a fraction of YTHDF2 targets are shared with ZNF121. Further, we observed that ZNF121 and YTHDF2 binding and localization on mRNA is highly correlative when plotting ZNF121 CITS around YTHDF2 CITS (and vice-versa) (Fig. 3F). When considering CITS, 3,937 (∼23%) of the 17,461 ZNF121 CITS in the CDS (Supplemental Fig. S3I), and 1,265 (∼35%) of the 3,566 CITS specifically in exons (Fig. 3G), overlap with YTHDF2 CITS within a 100 nt window. Therefore, we conclude that most ZNF121 mRNA transcript targets are also bound by YTHDF2, and that their RNA binding sites are coincident, i.e., within 100 nt of each other.

### The effect of ZNF121 on gene expression

To examine the effects of ZNF121 on gene expression, we utilized CRISPR/Cas9 to generate a ZNF121 knockout (KO) cell line and performed RNAseq (Supplemental Fig. S4A,B). The KO of ZNF121 was verified by Western blotting using an anti-ZNF121 antibody (Supplemental Fig. S4A). 1,110 genes and 1,318 genes were up- and down-regulated, respectively, in the ZNF121 KO cells relative to a WT-HEK293 control (Supplemental Table 4). Gene-set enrichment analysis enriched for terms including cellular homeostasis, RNA processing and transport, as well as terms related to cell death (Supplemental Fig. S4C). When comparing the proportions of up- and down-regulated genes from RNAseq with our iCLIPseq data and the ChIPseq data for ZNF121-bound promoters, ZNF121 slightly favoured binding to promoters over RNA in the differentially expressed genes. < 10% of regulated genes were bound by ZNF121 on both promoters and RNA. Among the differentially regulated genes bound by ZNF121 at the DNA or mRNA level, most were down-regulated when ZNF121 was knocked out, suggesting that ZNF121 may mostly up-regulate gene expression both as a transcription factor and as an RBP (Supplemental Fig. S4D).

RT-qPCR was performed to validate the effects of ZNF121 KO on the mRNA abundance for proteins that regulate the deposition and function of m6A. The ZNF121 KO had little effect for most of these genes, except PCIFI, a writer for the N^6^,2′-O-dimethyladenosine (m6Am) modification found at the first transcribed adenosine (Sendinc et al. 2019), which showed downregulation, whereas YTHDC1 and all three YTHDF paralogues were upregulated (Fig. 4A). These results were largely consistent with our RNAseq data (Supplemental Table 4,5). However, Western blot analysis showed no significant effect on the protein level of YTHDF2 (Supplemental Fig. S4E). It is possible that YTHDF2 protein synthesis or degradation is subject to negative feedback mechanisms, similar to the previously reported CDK1-YTHDF2-WEE1 loop (Fei et al. 2020) (Supplemental Fig. S4E).

**Figure 4.**
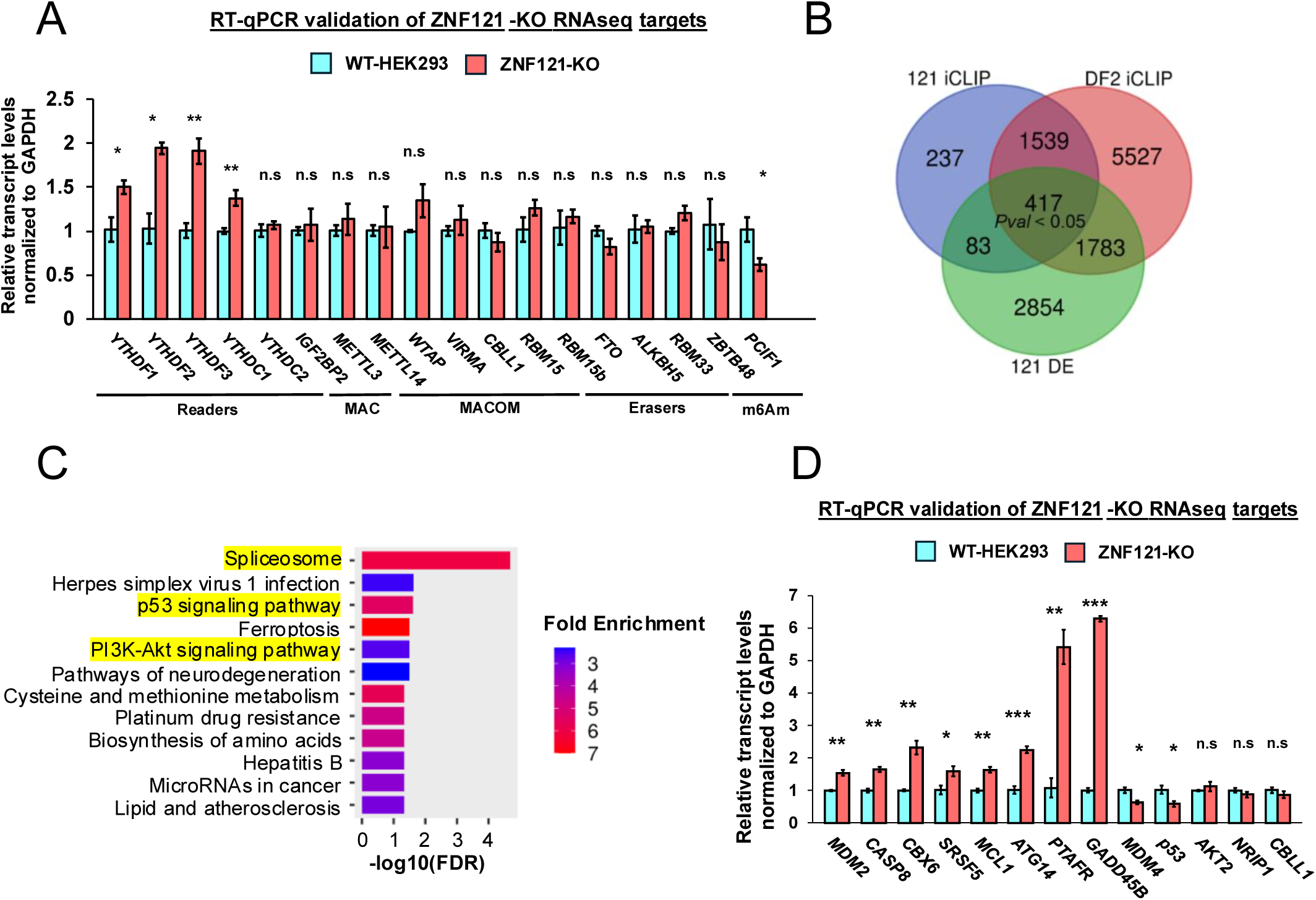
Knockout of ZNF121 results in changes to mRNA abundance of bound targets. (*A*) Validation of transcript abundance in the presence (blue) or absence (pink) of ZNF121 for transcript targets related to m6A biology. (*B*) Venn diagram showing overlap of transcripts containing ZNF121 CITS (121 CLIP), YTHDF2 CITS (DF2 CLIP), and those that are differentially expressed upon ZNF121 KO (121 DE). *p* derived from Hypergeometric Distribution. (*C*) KEGG enrichment analysis for the transcripts that have both ZNF121 and YTHDF2 CITS and are differentially expressed upon ZNF121 KO. (*D*) Validation of transcript abundance for a subset of the overlap targets identified in B that are related to splicing and the p53 and PI3K-Akt pathways. All qPCR data presented as ±SEM, n=3. *p,* ≤ 0.05 (*); ≤ 0.005 (**); ≤ 0.001 (***); > 0.05 (n.s) using two-tailed Student’s t-test.

We also determined the effect of deleting ZNF121 on targets bound by both ZNF121 and YTHDF2. The 417 overlapping YTHDF2∩ZNF121 differentially expressed targets upon ZNF121 KO (Log2Fold change cutoff ±0.05, *padj* ≤ 0.05) were enriched for terms including “spliceosome,” “p53 signalling pathways,” and “PI3K-AKT signalling pathways” in a KEGG pathway enrichment analysis (Fig. 4B,C). The transcript abundance for some of the genes implicated in the p53 pathway, i.e., *MDM2*, *MDM4*, *CASP8*, *p53,* and *GADD45B*, was also assessed by RT-qPCR in ZNF121 KO cells relative to WT-HEK293 cells. As expected, these analyses aligned with the RNAseq data, i.e., *MDM4* and *p53* were significantly downregulated whereas *MDM2*, *CASP8* and *GDD45B* were significantly upregulated (Fig. 4D, Supplemental Table 4).

### ZNF121 is important for the binding of YTHDF2 to shared targets

YTHDF2 was estimated to bind mRNA weakly, with K_d_ 2.5 µM and 21 µM in the presence or absence of m6A, respectively (Zhu et al. 2014). However, these *in vitro* experiments cannot account for other factors that might stabilize or destabilize the YTHDF2-mRNA interaction *in vivo*. To determine whether ZNF121 affects the binding of YTHDF2 to commonly bound targets, we performed RNA-immunoprecipitation (RIP) using YTHDF2 antibodies, followed by RT-qPCR. In these experiments, we observed that the ability of YTHDF2 to bind YTHDF2∩ZNF121 mRNA targets, but not mRNA targets bound only by YTHDF2, is impaired in ZNF121-depleted cells (Fig. 5A). These results suggested that ZNF121 plays a role in stabilizing the binding of YTHDF2 to shared targets.

**Figure 5.**
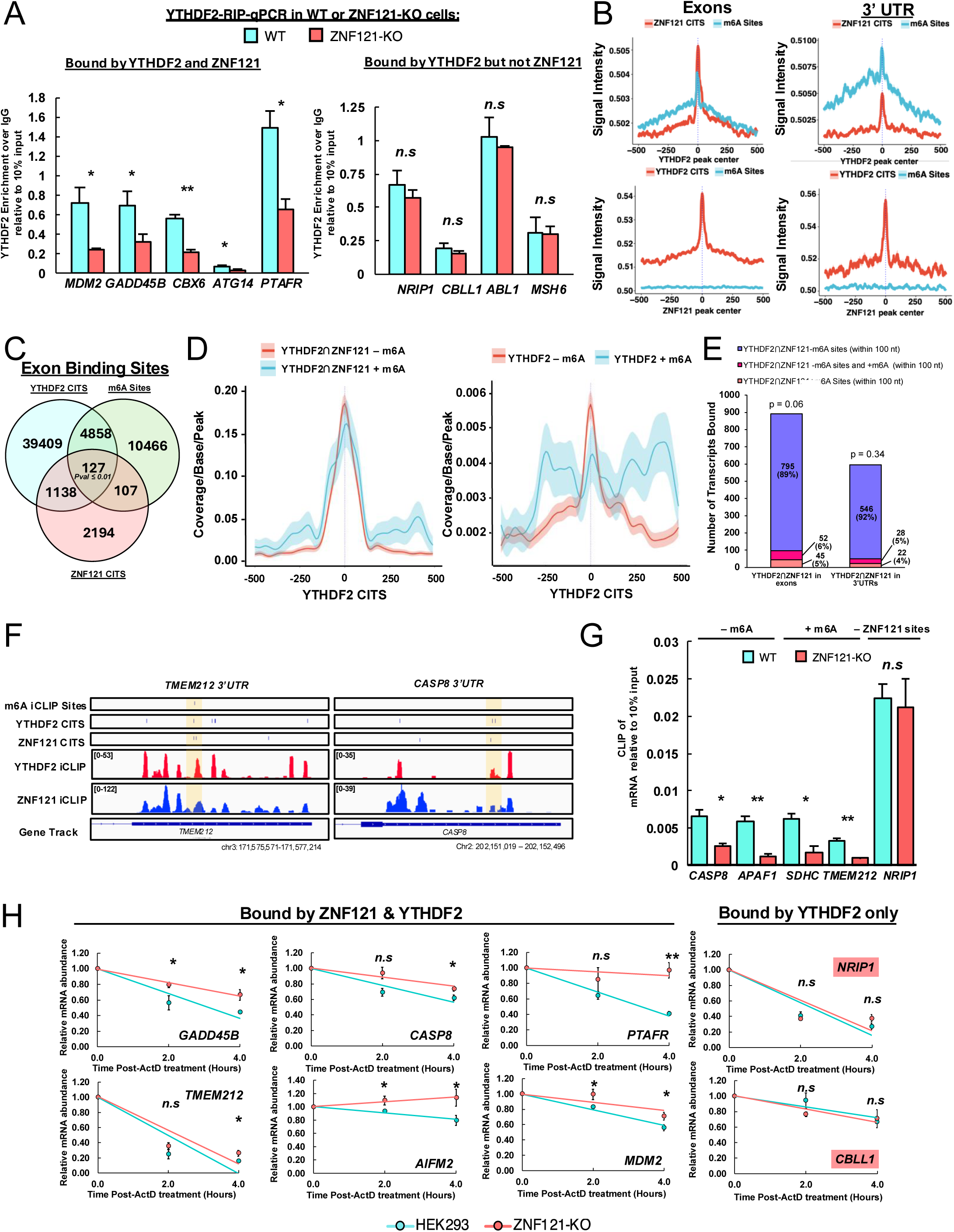
ZNF121 enhances the binding of YTHDF2 to common mRNA targets, independent of m6A, and controls their stability. (*A, Left*) YTHDF2 RIP-qPCR from extracts of ZNF121 KO or WT HEK293 cells of transcripts containing YTHDF2 and ZNF121 CITS, or (*right*) targets that have YTHDF2 CITS but no ZNF121 CITS. (*B*) Distribution of ZNF121 and YTHDF2 CITS and miCLIPseq m6A sites around YTHDF2 or ZNF121 sites, as indicated, either for all exons (*left*) or only for 3’UTRs (*right*). (*C*) Venn diagram showing the overlap of mRNA sites that contain ZNF121 CITS and/or YTHDF2 and/or m6A sites identified by miCLIPseq within 100 nt of each other. *p* determined by the Hypergeometric Distribution. (*D, Left*) Distribution of ZNF121 CITS around YTHDF2 CITS sites located near ZNF121 CITS within 100 nt in exons, or (*right*) ZNF121 CITS around all YTHDF2 CITS in exons, with or without nearby m6A (red and blue lines, respectively). (*E, left*) Numbers of transcripts, or (*right*) 3’UTRs containing nearby ZNF121 and YTHDF2 CITS, with or without nearby m6A sites identified by miCLIPseq (within 100 nt), as well as transcripts containing nearby ZNF121 and YTHDF2 sites with m6A sites either nearby or elsewhere in the transcript or 3’UTR. *p* determined by the Hypergeometric Distribution. (*F*) Example genome browser shots of transcripts with m6A iCLIPseq sites, YTHDF2 and ZNF121 CITS, and YTHDF2 and ZNF121 iCLIPseq reads (red and blue, respectively). Yellow shading indicates the 100 nt window around shared ZNF121 and YTHDF2 CITS. (*G*) YTHDF2 CLIP-qPCR in ZNF121 KO or WT-HEK293 cells for transcript targets containing YTHDF2 and ZNF121 sites, with or without miCLIPseq sites within 100 nt, as indicated. *NRIP1* is a negative control transcript bound by YTHDF2, but not ZNF121. (*H*) Decay of mRNA transcript targets of ZNF121 and YTHDF2 in ZNF121 KO or wild-type cells following actinomycin D (ActD) treatment. Red labels indicate genes which showed no change between conditions. All qPCR data presented as ±SEM, n=3. *p,* ≤ 0.05 (*); ≤ 0.005 (**); ≤ 0.001 (***); > 0.05 not significant (n.s) using Student t-test.

Because many of the mRNA binding sites of YTHDF2 co-localize with those of ZNF121 (Fig. 3F), we examined whether methylation status is related to their binding around common sites. We correlated the CITS of ZNF121 and YTHDF2 with our previously published m6A iCLIPseq (miCLIP) sites (Nabeel-Shah et al. 2024a, 2024b) in all exons or only those in 3’ UTRs (Fig. 5B). As expected, both ZNF121 and m6A sites are enriched around YTHDF2 binding sites (Fig 5B top). However, unlike YTHDF2, m6A sites are not enriched near ZNF121 binding sites on mRNA (Fig 5B bottom), suggesting that ZNF121 might stabilize the binding of YTHDF2 to RNA independently of whether YTHDF2 is bound near an m6A site.

To confirm this conjecture, we analyzed the relationship of ZNF121 and YTHDF2 binding sites to m6A sites within a 100 nt window. Only ∼ 10% of all YTHDF2∩ZNF121 binding sites within 100 nt of each other are near m6A sites (Fig. 5C). miCLIPseq sites do not enrich around YTHDF2∩ZNF121 sites located within 100 nt of each other and are depleted at ZNF121 sites that exclude YTHDF2 sites within 100 nt (Supplemental Fig. S5A). Plotting ZNF121 CITS around YTHDF2∩ZNF121 ± m6A located within 100 nt showed a correlation, regardless of m6A status, with a preference for non-methylated sites (ANOVA *p*= 7.89×10−^09^, 1.82×10^−21^ in exons or all features, respectively) (Fig 5D, left, Supplemental Fig. S5B). Plotting ZNF121 CITS around all YTHDF2 CITS ± m6A located within 100 nt showed that ZNF121 correlates well with YTHDF2 sites that exclude m6A, but less well with YTHDF2 sites having nearby m6A (ANOVA *p*= 3.2×10^−09^) (Fig. 5D right). These results showed that ZNF121 binds to mRNA near certain YTHDF2 sites regardless of whether m6A is present and that YTHDF2 itself can bind *in vivo* to certain sites on mRNA independent of m6A.

Of the 1,956 YTHDF2∩ZNF121 target transcripts (Fig. 3E), we found that 46% are bound by ZNF121 and YTHDF2 within 100 nt in all exons. Likewise, when considering the 1,190 targets bound in 3’UTRs, 52% are bound by YTHDF2 and ZNF121 located within 100 nt of each other (Supplemental Fig. S5C). Next, transcripts co-bound by YTHDF2 and ZNF121 were categorized based on the presence of nearby annotated m6A sites within 100 nt of the binding region. Among transcripts with YTHDF2∩ZNF121-bound exons, 795 (∼89%) lack nearby m6A sites, and among those with YTHDF2∩ZNF121-bound 3′ UTRs, 546 (∼92%) similarly lack nearby m6A sites. Only ∼5%-6% of YTHDF2∩ZNF121 co-bound transcripts contain shared binding sites in regions both with and without nearby m6A sites (Fig. 5E). These analyses further indicated that ZNF121 and YTHDF2 participate in mRNA binding independent of m6A.

To assess the robustness of m6A site identification, we compared our miCLIP-defined sites with published datasets generated using glyoxal and oxidative-reaction-based isothermal sequencing (GLORI), a non–antibody-based chemical method that enables identification and stoichiometric quantification of m6A sites (Liu et al. 2022). Approximately 80% of miCLIP sites are also detected by GLORI (Supplemental Fig. S5D). While many more m6A sites are identified by GLORI, our miCLIPseq analysis identifies the most frequently methylated sites (Supplemental Fig. S5E). YTHDF2 shares many RNA-binding sites with sites identified by both miCLIP and GLORI; however, most YTHDF2 sites are not near m6A sites in either analysis (Supplemental Fig. S5F,G). This is congruent with previous YTHDF2-PAR-CLIP results indicating ∼40% of YTHDF2 sites do not overlap with m6A (Wang et al. 2014a). Similarly, < 10% of ZNF121 mRNA binding sites are near any m6A site (Supplemental Fig. S5H,I). When overlapping ZNF121 and YTHDF2 CITS with GLORI m6A sites, ∼ 2.8% of all ZNF121 CITS overlap with YTHDF2 CITS, and of those, only ∼15% are near GLORI m6A sites (Supplemental Fig. S5J; Fig. 5C). Further, YTHDF2 CITS favour more frequently methylated sites, whereas the small number of ZNF121 sites near m6A sites favour low-frequency methylation sites (Supplemental Fig. S5K,L). Therefore, the overlap of ZNF121 CITS with YTHDF2 CITS tends to exclude m6A in both our miCLIPseq and GLORI analyses.

To further examine whether ZNF121 could facilitate m6A-independent binding of YTHDF2 to mRNA, YTHDF2 RIP-RT-qPCR from WT or ZNF121 KO HEK293 cells was performed. Targets that have few or no miCLIPseq sites and few YTHDF2 sites in the transcript (mostly restricted to the 3’UTR) within 100 nt of YTHDF2∩ZNF121 sites were examined (Supplemental Table 6). When considering both sets of transcript targets, the absence of ZNF121 impacts the binding of YTHDF2 to its targets regardless of m6A status within 100 nt (Supplemental Fig. S6A). For further validation, CLIP RT-qPCR was performed using UV-treated WT-HEK293 or ZNF121-KO cells. Primers were designed around sites bound by ZNF121 and YTHDF2 within 100 nt which have nearby m6A sites (*SDHC* and *TMEM212*), or do not have nearby m6A sites (*CASP8* and *APAF1*), or are bound by YTHDF2 but not ZNF121 (*NRIP1*). Genome browser shots illustrate some of these identified YTHDF2∩ZNF121 mRNA binding sites (Fig. 5F). As one example, the 3’UTR of *CASP8* shows YTHDF2∩ZNF121 iCLIPseq CITS within 100 nt of each other in the absence of any m6A site. In contrast, *TMEM212* shows YTHDF2∩ZNF121 CITS within 100 nt of an m6A site. The results confirmed that ZNF121 depletion significantly reduces YTHDF2 binding to their common transcript targets, regardless of m6A site status (Fig. 5G). Together, these observations suggest that ZNF121 stabilizes the binding of YTHDF2 to common mRNA targets, regardless of the local m6A status.

### ZNF121 regulates the stability of transcripts it binds together with YTHDF2

YTHDF2 facilitates target mRNA decay through the recruitment of the CCR4-NOT complex (Wang et al. 2014a; Du et al. 2016). Therefore, we measured the stability of some of the target transcripts bound by YTHDF2 and affected by the presence of ZNF121. WT or ZNF121 KO HEK293 cells were treated with Actinomycin D (ActD) for 0h, 2h, or 4h, and the abundance of target transcripts was measured at each time point (Fig. 5H). The absence of ZNF121 imparted a significant stabilizing effect on transcripts upregulated upon ZNF121 KO (Fig 4D, Supplemental Table 4; Log2FoldChange ≥ 0.5, *padj* ≤ 0.05) and bound by ZNF121 and YTHDF2, such as *CASP8* and *MDM2* (Supplemental Table 4). In comparison, transcripts bound by YTHDF2, but not ZNF121, showed no significant difference in stability (Fig. 5H). This indicated that ZNF121 could be affecting mRNA stability through YTHDF2.

Since ZNF121 has a negative effect on the half-life of target transcripts also bound by YTHDF2, and YTHDF2 negatively affects the length of poly(A) tails by recruiting the CCR-NOT complex (Wang et al. 2014a; Du et al. 2016), we also tested the effect of ZNF121 on Poly(A) tail lengths. In comparison to the WT, a subtle lengthening of the poly(A) tails for *MDM2* and *CASP8* is observed in the ZNF121 KO (Supplemental Fig. S6B). In contrast, *CBLL1*, which is bound by YTHDF2 but not ZNF121, shows minimal change in average tail length.

Luo et al. (2018) found in a yeast two-hybrid screen that ZNF121 interacts with the PAN2 subunit of the PAN2-PAN3 deadenylase complex. Consistent with that, YTHDF2 and PAN2 co-purify with GFP-ZNF121 (Supplemental Fig. S6C), but CNOT1 does not. Moreover, ZNF121 and CNOT1, but not PAN2, co-purify with YTHDF2, as reported (Du et al. 2016) (Supplemental Fig. S6D). These results indicated that ZNF121 may influence target transcript stability and the rate of poly(A)-tail decay not only by enhancing the RNA-binding of YTHDF2 but also, potentially, by its interaction with PAN2.

### ZNF121-mediated mRNA decay of MDM2 affects the DNA damage response and cell viability

We found that, when considering the 1,190 genes whose 3’ UTRs are bound by both ZNF121 and YTHDF2, p53 signalling, cell cycle, and cellular senescence are enriched in KEGG pathways (Supplemental Fig. S7A,B), and we showed that many of these transcripts have less binding by YTHDF2 and increased stability in the absence of ZNF121 (Fig. 5A,G-H, Supplemental Fig. S6A,B). To identify a potential biological consequence of mRNA binding by ZNF121, we investigated the effects of ZNF121 on *MDM2*, a key player in the cell cycle and DNA damage responses (Reza Saadatzadeh et al. 2017). Consistent with our ZNF121 KO RNAseq data (Supplementary Table 4; Log2Fold change ≥ 0.05, *padj* ≤ 0.05), RT-qPCR showed that ZNF121 KO significantly upregulates *MDM2* (Fig. 4D), whereas overexpression of GFP-ZNF121 significantly reduces *MDM2* transcript abundance (Fig. 6A). Moreover, the depletion of ZNF121 significantly increases MDM2 protein abundance (Fig. 6B). These data highlight the consequences of the destabilizing effect of ZNF121 on *MDM2* mRNA (Fig. 5H), which results in lowered MDM2 protein abundance. Interestingly, the depletion of PAN2-PAN3 or CCR4-NOT deadenylase core components, as well as the KO of YTHDF2 itself, do not affect *MDM2* transcript abundance, suggesting ZNF121 is particularly important for the degradation of *MDM2* mRNA (Supplemental Fig. S7C-F).

**Figure 6.**
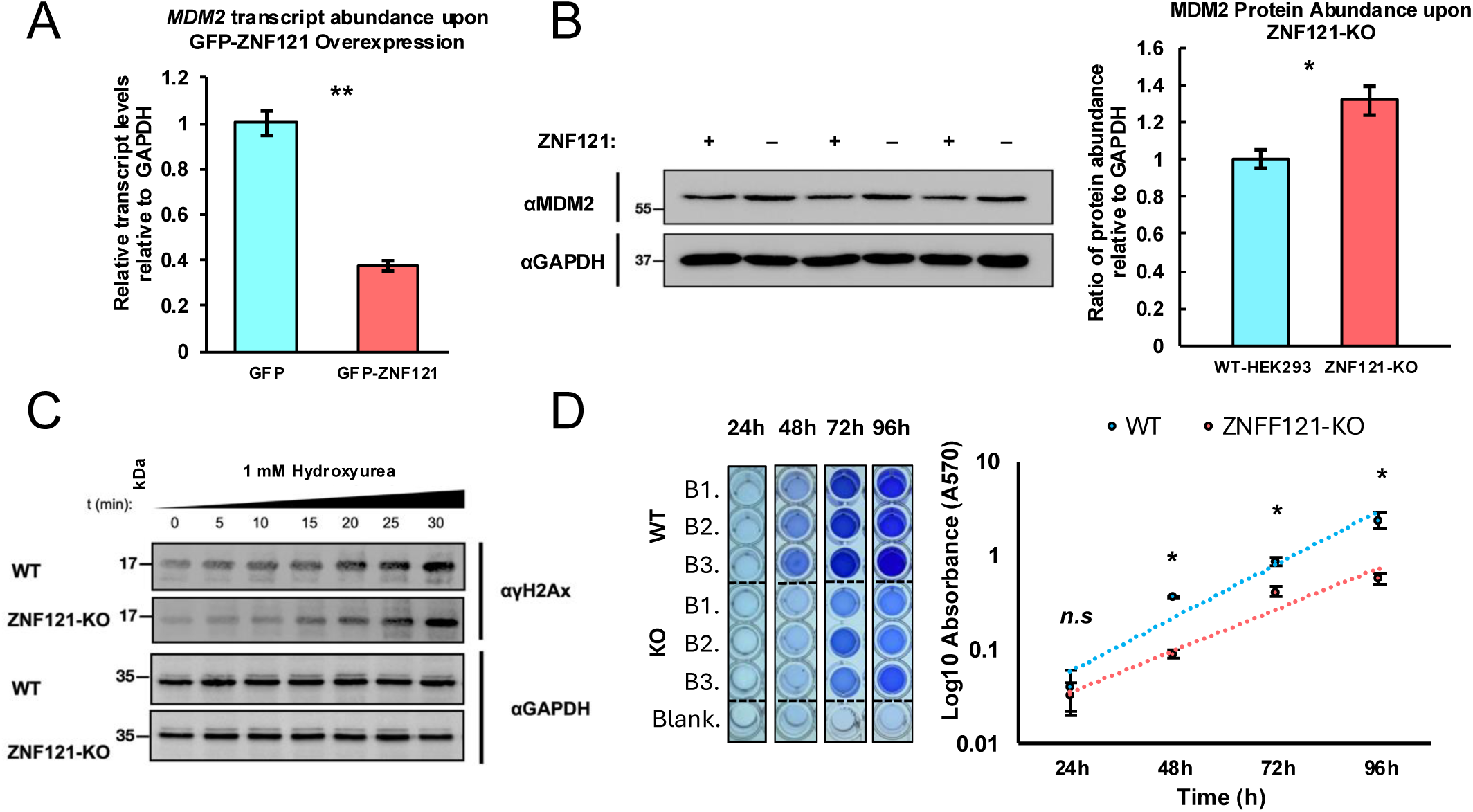
Knockout of ZNF121 results in increased MDM2 protein abundance, delayed DNA damage response and slower cell growth. (*A*) *MDM2* transcript levels based on RT-qPCR analysis in HEK293 cells over-expressing GFP-ZNF121, normalized to the levels in cells over-expressing GFP. (*B, left*) Western blot of MDM2 and GAPDH in the presence or absence of ZNF121, and (*right*) the corresponding quantification. (*C*) Western blots showing H2A.X activation in lysates from WT or ZNF121 KO HEK293 cells following treatment with 1 mM HU for the indicated times. (*D, Left*) Crystal Violet Assay, and (*right*) the quantification of ZNF121 KO or WT HEK293 cell growth. Wells were initially seeded with 6×10^3^ cells/well. Data presented as ±SEM, n=3. *p,* ≤ 0.05 (*); ≤ 0.005 (**); ≤ 0.001 (***); > 0.05 not significant (n.s) using Student t-test.

MDM2 is an E3-ubiquitin ligase and oncoprotein that binds and ubiquitinates p53 to maintain low, steady state levels of p53 in normal conditions. Upon DNA damage, MDM2 releases p53, facilitating activation by p53 of its downstream transcriptional program (Reza Saadatzadeh et al. 2017). While ZNF121 KO affects *p53* transcript abundance (Fig. 4D), YTHDF2 binding to *p53* mRNA (Supplemental Fig. S6A), and MDM2 protein levels (Fig. 6B), it does not alter p53 protein abundance or the mRNA levels of its downstream transcriptional targets *p21* and *p27* (Supplemental Fig. S7G-H). Therefore, the subtle regulation of MDM2 by ZNF121 might not be sufficient to derail the regulation of p53.

MDM2 also maintains the fidelity of the DNA damage response (DDR), independent of p53, by inhibiting the activation of H2A.X. H2A.X is a variant of histone H2A that is phosphorylated by the protein kinase ATM at Ser139 to form γH2A.X, which signals for the recruitment of the DDR machinery (Bouska et al. 2008; Alt et al. 2005). Given the above results, i.e., ZNF121-mediated regulation of MDM2 not in the context of the p53 pathway, we therefore tested for the potential role of ZNF121 in the DDR pathway. To this end, we investigated H2A.X activation in WT and ZNF121 KO HEK293 cells. Cells were left either untreated or treated with 1 mM hydroxyurea (HU) over 24h to induce DNA damage. A non-significant decrease in γH2A.X signal intensity was observed for cells exposed to HU for 24h, but untreated ZNF121 KO cells showed a ∼72% decrease in H2A.X activation compared to WT (Supplemental Fig. S7I). We then repeated this experiment with samples taken every 5 min for 30 min after treatment with HU – the determined optimal window for detecting changes in H2A.X activation (Supplemental Fig. 8A) – we again saw less initial γH2A.X in ZNF121 KO cells compared to WT. Further, there was a delay in γH2A.X signal over time in ZNF121 KO cells until ∼30 min, when the signal from ZNF121-KO cells was ∼80% that of WT (Fig 6C, Supplemental Fig. 8B). Importantly, analysis of our ZNF121 KO RNAseq data shows that ZNF121 does not affect H2A.X expression (Supplemental Table 4), making it unlikely that ZNF121 transcriptionally affects H2A.X abundance. These results showed that ZNF121 affects the activation of H2A.X upon DNA damage, consistent with the effect of ZNF121 on MDM2 expression and the known function of MDM2 in the DDR (Bouska et al. 2008; Alt et al. 2005).

Previous literature indicates that the depletion of ZNF121 using siRNA results in reduced cell growth in cancer cell lines (Luo et al. 2016). To investigate this further, we utilized the Crystal Violet Cell Viability assay on ZNF121 KO HEK293 cells. Consistent with earlier findings, ZNF121 KO cells exhibited slower growth compared to WT cells (Fig. 6D, Supplemental Fig. S8C). ZNF121 was also implicated in cell cycle regulation and breast cancer progression, correlating with reduced survival in patients (Luo et al. 2018, 2016). We show that ZNF121 and YTHDF2 interact in MCF-7 breast cancer, as well as HCT-116 colorectal adenocarcinoma cells (Fig. 1A). Notably, ZNF121 is significantly overexpressed across all three subtypes of colorectal cancer (Supplemental Fig. S8D), and patients with tumors expressing high levels of ZNF121 initially experience poor outcomes compared to those with relatively low ZNF121 levels (Supplemental Fig. S8E). These observations suggest a potential role for ZNF121 in colorectal cancer, in addition to its previously reported role in breast cancer (Luo et al. 2018, 2016).

## Discussion

We have shown here that ZNF121 interacts with YTHDF2 in the cytoplasm, binds to mRNA near YTHDF2, and stimulates the interaction of YTHDF2 with their common mRNA targets. We discovered that most ZNF121-YTHDF2 common RNA binding sites lack m6A marks. Further, deleting ZNF121 weakens YTHDF2’s binding to these targets, resulting in increased transcript stability. In this manner, we have identified a novel m6A-independent mechanism for YTHDF2-RNA binding, with downstream effects on target mRNA stability.

YTHDF2 utilizes residues outside its tryptophan cage to bind the phosphate backbone of RNA and uses its tryptophan cage to recognize m6A in a two-step process. *In vitro* studies showed that the ablation of the m6A-recognizing tryptophan cage diminishes but does not eliminate RNA-binding by YTH-proteins. However, mutating residues which interact with the RNA phosphate backbone eliminated RNA-binding (Zhu et al. 2014). That is presumably why previous CLIP-based experiments identified YTHDF2-binding sites that do not overlap with identified m6A sites (Shi et al. 2017; Wang et al. 2015). Similarly, our iCLIPseq analyses indicated that many YTHDF2 crosslink sites on mRNA are not located near m6A sites identified by either miCLIPseq or GLORI.

While we found that ZNF121 does stimulate the binding of YTHDF2 to some sites containing m6A (consistent with our observation that the RNA and m6A binding sites of YTHDF2 remain open in the YTHDF2-ZNF121 heterodimer structure predicted by AlphaFold-Multimer), ZNF121 stimulates the binding of YTHDF2 to many more sites that are not methylated. Since most YTHDF2 crosslinking sites are not shared with ZNF121, there may be other proteins, perhaps many of them, that interact with YTHDF2 to recruit it to specific sets of RNA binding sites. One example, HRSP12, engages in cooperative binding with m6A-bound YTHDF2 on mRNAs that also have HRSP12 binding sites, facilitating the recruitment of the RNaseP/MRP complex for target endonucleolytic cleavage (Park et al. 2019).

YTH-RNA interactions may also be stabilized by biomolecular condensate formation, since YTHDF2 undergoes biomolecular condensation upon binding to methylated mRNA *in vitro* (Ries et al. 2019). Further, YTHDF2 localizes to P-bodies, phase-separated cytoplasmic foci that facilitate mRNA processing, storage, and decay and promote stress granule formation (Wang et al. 2014a). The formation of these granules, or cooperative binding with RBPs, might reduce transient binding between YTHDF2 and its targets, stabilizing the RNA-protein complex.

YTHDF2 plays a key role in poly(A) tail decay and mRNA destabilization (Wang et al. 2014a; Du et al. 2016). Deadenylation is believed to be biphasic: first, PABPC1 bound to poly(A)-tails recruits the PAN2-PAN3 complex to slowly initiate poly(A) decay; second, the CCR4-NOT complex is recruited when poly(A)-tails are reduced to approximately 100 nt in mammalian cells, rapidly completing decay and promoting further mRNA processing (Passmore and Coller 2021). We confirmed that ZNF121 interacts with PAN2 in co-IP experiments, but not CNOT1, whereas YTHDF2 interacts with CNOT1, but not PAN2, in similar experiments, reaffirming previous observations (Luo et al. 2018; Du et al. 2016). Based on these findings, ZNF121 may bind to target mRNAs and recruit PAN2 to initiate poly(A)-tail decay and stabilize the YTHDF2-mRNA interaction, enabling more efficient recruitment of CCR4-NOT, consistent with the biphasic deadenylation model. Indeed, depleting cells of YTHDF2, PAN2, or CNOT1 alone does not affect *MDM2* abundance. Only when ZNF121 is deleted did we observe increased *MDM2* mRNA abundance. Thus, ZNF121 may prime certain YTHDF2-bound transcripts for efficient poly(A)-tail shortening and mRNA clearance by recruiting PAN2 and functioning in concert with YTHDF2. In this respect, ZNF121 may be analogous to ZFP36 and other TTP family members that bind AU-rich elements (AREs), Pumilio-family proteins that bind the Pumilio-response element (PRE), and YTHDF2 itself, all of which facilitate targeted recruitment of CCR4-NOT (Van Etten et al. 2012; Lykke-Andersen and Wagner 2005; Passmore and Coller 2021; Du et al. 2016). This potential role of ZNF121 in biphasic poly(A)-tail decay and mRNA destabilization warrants further exploration, particularly regarding its ability to interact with, and recruit, PAN2 to mRNA.

We identified ZNF121 as another C2H2-ZFP RBP and used AlphaFold 2.0 to predict its right-handed helical structure. This predicted helical structure is also true for the ZnF regions of C2H2-ZFPs whose structures have been determined (Pavletich and Pabo 1991; Yang et al. 2023; Pavletich and Pabo 1993; Houbaviy et al. 1996). The tandem ZnFs of C2H2-ZFPs wind around target DNA (Fedotova et al. 2017), inserting their DNA-recognizing α-helices into the major groove. There was no statistically significant overlap between the DNA and RNA sites bound by ZNF121. We also found that no specific set of several tandem ZnFs of ZNF121 is essential for the interaction of ZNF121 with RNA, raising the possibility that different sets of ZnFs may recognize different sequences or structures in RNA, as can be the case for DNA-binding by the C2H2-ZFPs (Jolma et al. 2024). Whether different subsets of the ZnFs of ZNF121 bind distinct RNAs remains unclear. The IDR of ZNF121 may also contribute to RNA binding, consistent with recent reports indicating C2H2-ZFPs IDRs involvement in RNA binding (Gosztyla et al. 2024).

We found that ZNF121 deletion leads to slower growth of HEK293 cells, consistent with previous studies on T-47D, MDA-MB-231, and MCF10A breast cancer cells using siRNAs against ZNF121 and the observation that ZNF121-depleted cells remain in G_1_/G_0_. (Luo et al. 2016). However, whereas ZNF121 depletion did not affect p53 abundance in our ZNF121 KO system, Luo et al. (2016) reported that ZNF121 overexpression decreased the protein abundance of p53 and increased those of its downstream targets *p21* and *p27*. This discrepancy might reflect differences in cell types used in these experiments or from comparing the results of a gene knockout with those obtained using protein overexpression. In addition, the regulation of p53 is layered and complex (Liu et al. 2024).

Luo et al. (2016, 2018) reported that ZNF121 is overexpressed in breast cancer tumours and regulates MYC and BRCA1. We similarly found that ZNF121 is also significantly overexpressed in all three subtypes of colorectal cancer and identified the interaction of ZNF121 and YTHDF2 in HCT-116 colorectal carcinoma cells. Additionally, patients whose tumors express high levels of ZNF121 exhibit poorer outcomes. Among the genes affected by ZNF121, cell cycle regulators were significantly enriched, including *MDM2*, a key regulator of the DNA damage response. Through its interaction with YTHDF2, ZNF121 stabilizes YTHDF2’s binding to target mRNAs and promotes their decay. Loss of ZNF121 likely alters the expression of other cell cycle genes, contributing to the impaired cell fitness observed upon ZNF121 depletion. The ZNF121-YTHDF2 interaction may therefore play a role in ZNF121s pathological associations with breast and colon cancer.

## Materials and Methods

### Cell culture

HEK293 Flp-In™T-Rex cells (Invitrogen, Cat# R780-07), HCT-116 and MCF-7 cells from ATCC (CCL-247 and CRL1573, respectively), and HeLa cells (kindly donated by Dr. Jason Moffat) were maintained as described in (Nabeel-Shah et al. 2022).

#### Generation of stable cell lines

Gateway^TM^-compatible ORFs were inserted into the pDEST pcDNA5/FRT/TO-eGFP vector according to the manufacturer’s instructions (Invitrogen™, Cat# 11789-013). The vector was co-transfected into HEK293 Flp-In™T-REx cells with the pOG44 Flp recombinase expression plasmid, and individual clones were selected with hygromycin (Life Technologies, 10687010) at 200 μg/mL for FRT site-specific recombination into the genome. Expression was induced with Doxycycline (1 μg/mL) in the culture medium 24 hours prior to cell harvesting and validated by western blot analysis. Lipofectamine® 2000 (Invitrogen™, Cat# 11668019), Opti-MEM™ Reduced Serum Medium (Gibco, Cat# 31985070) and 3 μg of plasmid DNA per replica were used for transient transfections over 48h.

### siRNA-mediated knockdowns

Silencer Select experimental and control siRNAs from Thermo Fisher Scientific (METTL3: Cat # s32142, s32143; PAN2: Cat # s19252, s19253; CNOT1: s22842, s22843; control non-targeting siRNA: AM4611) were delivered into cells with Lipofectamine® RNAiMAX Reagent (Invitrogen™, Cat# 13778-075) in biological triplicates for 72h.

### Cloning and plasmid construction

The Gateway^TM^ cloning system was used to clone Accuprime PFX DNA Polymerase (Thermo Fisher Scientific, CAT# 123440024) PCR products containing flanking ATTB sites (5 μg) into pDONR223 vectors, in TE buffer pH 8.0 using Gateway^®^ BP Clonase Enzyme Mix (Invitrogen™, Cat# 11789-013) for 2h to overnight, followed by Proteinase K digestion (Invitrogen™, Cat# EO0491) at 37°C for 10 min, then incubation on ice for 3 min.

For expression vector transformation, the BP Clonase mixture was incubated with 50 μl of competent *DH5*α competent *E. coli* cells (Invitrogen™, Cat# 18265017) on ice for 20 min, followed by 25s heat shock at 42°C, 5 min ice incubation, supplementation with 950 μl of LB liquid media and 1h incubation, shaking at 37°C and 250 RPM. Cells were briefly centrifuged at 8000x g, resuspended and spread on LB solid agar plates with 100 μg/ml spectinomycin and grown overnight at 37 °C. Colonies were incubated overnight in liquid LB containing 100 μg/ml spectinomycin at 250 RPM and 37 °C. The Presto™ Mini Plasmid Kit was used to extract plasmids (Geneaid™, Cat # DFH300) which were incubated with pDEST pcDNA5/FRT/TO containing 5’GFP or 5’ FLAG-tags and LR Clonase Enzyme Mix (Invitrogen™, Cat# 11791019), followed by transformation as described above, using 100 μg/ml ampicillin for selection.

Q5® Site-Directed Mutagenesis was performed according to manufacturer’s recommendations (New England Biolabs®, Cat# E0554S), using https://nebasechanger.neb.com/ for primer design.

See Supplemental Table 7 for a complete list of primers used.

### CRISPR-mediated gene knockouts

CRISPR-mediated gene knockouts were performed using the LentiCRISPRv2 plasmid, which was kindly provided by the Moffat Lab (Addgene plasmid #52961). Guide RNA (gRNA) sequences from the Toronto KnockOut version 3 (TKOv3) library for ZNF121 (AATGTGGAAGAGCCTTCGCT) and YTHDF2 (ATATAGGTCAGCCAACCCAG) were each cloned into the LentiCRISPRv2 vector and transfected into 293T cells to produce lentiviral particles Flp-In™T-REx HEK293 cells were infected with lentivirus at an MOI of ∼0.3 in the presence of polybrene (8ug/mL; Sigma-Aldrich, Cat # H9268), selected with puromycin (2μg/mL; BioShop Canada Inc., Cat # PUR333.25) for 3 days and grown for 3 more days. Single cells were sorted into 96-well plates containing DMEM (20% FBS and 10% conditioned medium). Plates were checked for growth of single cell colonies after one week, and clonal populations were expanded and knockouts confirmed by Western blotting.

### Western Blot

Samples were suspended in sample buffer (140 mM Tris-HCL, pH 6.8, 4% SDS, 20% glycerol and 0.02% bromophenol blue) with 1:100 2-mercaptoethanol (BioRad, Cat# 1610710XTU), boiled for 5 min, briefly centrifuged, followed by SDS-PAGE (90v – 180v) in SDS,Tris and Glycine buffer. Proteins were transferred to methanol-activated PVDF membranes (Caledon Laboratory Chemicals, Cat# 67601-7-40, BioRad, Cat# 1620177) at 100v for 1h in transfer buffer (50 mM Tris pH 7.5, 380 mM Glycine, 20% v/v methanol) and subsequently blocked in 1xTBST (0.05 M Tris, 0.15 M NaCl, 0.1% Tween-20, at a of pH 6.7), 5% w/v skim milk (BioShop Canada Inc., Cat# SKI400.500) for 45 min, then washed with 1xTBST and incubated with the corresponding primary antibody diluted in 0.25% w/v Bovine Serum Albumin (BioShop Canada Inc., Cat# ALB005.250) at 4°C overnight. Membranes were washed for 5 min, three times in 1xTBST, incubated with 1:5000 secondary antibody in blocking solution at room temperature and washed again for 5 min, three times. ECL Western Blotting Detection Reagents (Cytiva™, Cat# RPN2106-OL-AB) was used for signal detection in the DNR BioSystems MicroChemi 4.2. Quantification was performed on three biological replicates using ImageJ (Abràmoff et al. 2024).

Antibodies are listed in Supplemental Table 8.

### Immunoprecipitation

IPs were performed as previously described (Nabeel-Shah et al. 2024a, 2024b). At ∼80% confluency, cells were resuspended in ice-cold IP lysis buffer (10 mM Tris pH 7.5, 140 mM NaCl, 1 mM EDTA, 1% Triton-X100, 0.1% Sodium Deoxycholate) supplemented with protease inhibitors (Roche, Cat# 05892791001), sonicated (0.3sec on, 0.7 sec off, amplitude of 30%) and treated with 75 units of Benzonase Nuclease (Sigma-Aldrich, Cat# E8263-25KU) for 30 min on a rotator at 4°C. Lysates were clarified at 15,000x g for 30 min. 1% of each lysate was retained as an input. The rest of the lysates were incubated with Dynabeads Protein G magnetic beads (Invitrogen™, Cat# 10003D) pre-conjugated with the appropriate antibodies, as listed in Supplemental Table 8, and rotated overnight at 4°C. Fresh beads were washed with the IP lysis buffer and pre-conjugated with the relevant antibodies for 2h, rotating at 4°C. Beads were washed with the IP lysis buffer (with 2% TritonX-100 and 1% NP40) for 5 min three times at 4°C and resuspended in 1x SDS sample buffer (as with inputs) containing 1:100 2-mercaptoethanol (BioRad, Cat# 1610710XTU), followed by western blot.

### Subcellular fractionation

This assay was adapted from Morcos et al. (2025). Cells grown in 15 cm cell culture plates were harvested with 2 ml of 0.05 % trypsin and 0.53 mM EDTA (Winset Inc., Cat # 325-542-CL), resuspended in ice-cold 1xPBS (Winset Inc., Cat# 311-010-LL), and pelleted at 800x g for 5 min. Pellets were washed with 5 pellet volumes of ice-cold 1xPBS, centrifuged at 800x g for 5 min and resuspended in two pellet volumes of cytosolic lysis buffer (20 mM Tris pH7.7, 10 mM NaCl, 1mM EDTA, and 0.5% NP40) supplemented with protease inhibitors (Roche Cat# 05892791001). The suspension was centrifuged at 500x g for 5 min at 4°C, and the supernatant cytoplasmic fraction was retained and centrifuged briefly. The nuclei-containing pellet was washed three times with two pellet volumes of neutral buffer (20 mM Tris pH7.7, 0.2 mM EDTA, 20% glycerol v/v), centrifuged at 4°C for 5 min, and resuspended in three volumes of nuclear extraction buffer (20 mM Tris pH7.7, 300 mM NaCl, 1mM EDTA, and 20% glycerol v/v) supplemented with protease inhibitors (Roche, Cat# 05892791001), and centrifuged at 15,000x, 4°C for 15 min. Both fractions were sonicated and clarified at 15,000x g for 30 min at 4°C. ∼ 30 µg of protein were mixed with SDS gel sample buffer supplemented with 1:100 2-mercaptoethanol (BioRad, Cat# 1610710XTU) and subjected to western blot.

For IP, both fractions were adjusted to 150 mM NaCl, followed by the IP protocol as above.

### Immunofluorescence

Cells seeded on Poly-L-Lysine-coated microscope cover slips (Sigma-Aldrich, Cat# P8920). At 70% confluency, cells were induced with 1μg/ml of Doxycycline (Sigma-Aldrich, Cat# D9891) for 24h, fixed for 20 min with 1% PFA (Electron Microscopy Sciences, Cat#15710) w/v in 1xPBS (Winset Inc. Inc, Cat# 311-010-LL) and 0.02% TritonX-100 (Fluka BioChemika, Cat#93418) at a pH of 7.5, washed three times for 5 min with 1xPBS and permeabilized with 0.5% Tergitol solution (Sigma-Aldrich, Cat# NP40S-10ML) for 10 min at room temperature, then washed with 1x PBS three times for 5 min. Cells were blocked for 60 min at room temperature with blocking buffer containing 1 % New Goat Serum (NGS) (Rockland, Cat# B304) and 3% bovine serum albumin (BSA, Winset Inc., Cat# 800-0950CG) in 1x PBS. Cover slips were incubated overnight in a humidity chamber with antibodies (Supplemental Table 8) diluted in blocking buffer. Cover slips were washed three times for 5 min each with 1xPBS and incubated with the corresponding secondary antibody (Rockland, Cat# 610-152-121S, 611-156-122S) diluted in blocking buffer (1:250) in a humidity chamber, then washed 3 times for 5 min with 1xPBS, incubated with 1:1000 DAPI (4’,6-diamidino-2-phenylindole) stain (Sigma-Aldrich, Cat# D9542-1MG) in 1xPBS for 10 min, washed once with 1x PBS, mounted on microscope slides, cured overnight and stored at 4°C. Images were acquired using a spinning disk confocal microscope at the imaging facility of the PGCRL, SickKids, Toronto, Canada.

### RNA stability assays

These assays were performed as described previously (Nabeel-Shah et al. 2022). Cells were collected at ∼80% confluency, seeded at a density of 0.3×10^6^ cells per-well in 6-well culture plates, and incubated at 37°C until ∼80% confluent. Cells were treated with 0.01 mg/ml of Actinomycin D (BioShop Canada Inc., Cat# ACT001.5) suspended in 1xPBS (Winset Inc. Cat# 311-010-LL) and incubated for 0h, 2h, or 4h at 37 °C, harvested by trypsinization (Winset Inc. Cat# 325-542-CL) and collected in ice-cold 1xPBS. *HPRT1* was used as a loading control in RT-qPCR analysis, F: CATTATGCTGAGGATTTGGAAAGG, R: CTTGAGCACACAGAGGGCTACA.

### RNA extraction and cDNA synthesis

RNA was extracted with 1 ml of TRIzol Reagent (Invitrogen™, Cat# 15596026), per manufacturer’s instructions, and cDNA was then prepared using the SuperScript® VILO™ cDNA Synthesis Kit (Invitrogen™, Cat# 11754-050).

### RT-qPCR

PowerUp™ SYBR Green Master Mix (Applied Biosystems™™, Cat# A25742) was mixed with cDNA and primers, and run on the Applied Biosystems™ 7300 real time PCR System (Thermo Fisher, Cat# 4406984) programmed to 40 cycles of 95 °C for 15 s, 60 °C for 30 s, and then 95 °C for 15 s and 60 °C in three biological replicates, and at least three technical replicates. The relative transcript abundance was calculated using the ΔΔCT method (Livak and Schmittgen 2001) relative to GAPDH or HPRT1, and to t = 0h for stability assays. qPCR analysis utilized the threshold cycles (CTs) generated by QuantStudio^TM^ Real-Time PCR to calculate expression levels. Primers used for RT-qPCR are available in Supplemental Table 9.

### RIP RT-qPCR

Cells were harvested at ∼75% confluency (n=3), resuspended in 1 mL of ice-cold RIP lysis buffer (150 mM NaCl, 25 mM Tris pH 7.5, 0.5% NP 40, and DEPC treated water) and supplemented with protease inhibitors (Roche, Cat# 05892791001). 4000 u/mL of RNase Inhibitor Murine (New England Biolabs®, Cat# M0314L) was added and samples were incubated on ice for 30 min, and clarified at 15,000x g for 20 min at 4°C retaining the supernatant. 10% was set aside for RNA input, and the remainder was incubated with pre-conjugated beads. 50 μl Dynabeads Protein G (Invitrogen™™, Cat# 10003D) were washed briefly three times with RIP lysis buffer and conjugated to 5 μg of anti-YTDHF2 antibody (Proteintech, Cat# 24744-1-AP) or rabbit-IgG antibody (Invitrogen™, Cat# 10500C) for two hours of end-to-end rotation in 100 μl/sample of RIP lysis buffer at 4°C. Beads were washed three times with 1 ml ice cold RIP wash buffer (150 mM NaCl, 25 mM Tris pH 7.5, 0.05% NP-40 and DEPC treated water), then suspended in 1ml of TRIzol reagent for RNA extraction (Invitrogen™™, Cat# 15596026). 5-10 μg of glycogen (Invitrogen™, Cat# R0551) was added as the co-precipitant with RNA, which was resuspended in 5 μl UltraPure dH_2_O (Invitrogen™, Cat# 10977-015) and quantified for RT-qPCR.

Analysis was relative to 10% of input RNA, and IgG signal from the RT-qPCR was subtracted from the YTHDF2-IP.

### Poly(A) tail length assay

Poly(A) tail length assays were performed according to the manufacturer’s instructions (Thermo Scientific™, Cat # 764551KT). The primers used here are as follows: *MDM2 F CCTTTCGTTTGTTAGCTCATTT, R* CTACCATGTAGCCAGCTTTC; CASP8 F ACTTGCTTTATGCCTTCTTATTG, R AATCCAAGCAGAGATGAAAGA; CBLL1 F GAGGCATATCAGATGAATGTT, R TAAGAATTAAACTGCCACCTC. Universal reverse primers were included in the kit.

### Crystal violet assays of cell viability

Cells were grown to ∼ 80% confluence, seeded at 6×10^3^ cells per-well in 96-well plates (n=3) and incubated at 37°C and 5.0% CO_2_. At the indicated times, Life Technologies EVOS FL (20x) was used for microscopy and cells were processed using the Crystal Violet Cell Viability Assay (Abcam, Cat# AB232855) kit per the manufacturer’s instructions. Opacity of the wells was photographed on a backlight, quantified using a BioTek EPOCH Multiplate Spectrophotometer and BioTek Gen5 (v2.01). Cell viability was calculated per the Crystal Violet Cell Viability Assay instructions.

### H2A.X activation assay

Cells were grown to ∼ 80% confluency, and 3.2×10^6^ cells were seeded into 60 mm culture plates. At ∼80% confluency, the media were changed to including 1 mM hydroxyurea (BioShop Canada Inc., Cat# HYD023.10) and allowed to incubate at 37°C, 5.0 % CO_2_ for the indicated times. Cells were harvested by trypsinization, resuspended in ice cold 1xPBS, and centrifuged at 4000x g for 5 min and subject to Western blot analysis. Anti-γH2A.X (Abcam, Cat# ab81299) or GAPDH (Invitrogen, Cat# 39-8600) as a loading control were used. ImageJ was used to quantify the signal.

### RNA sequencing

Total RNA was extracted from wild-type or ZNF121-KO HEK293 cells in biological duplicates. RNA was extracted from cell pellets using TRIzol reagent (Invitrogen™, Cat# 15596026), and DNA was removed using DNase treatment followed by cleanup using the Zymo Research RNA Clean and Concentrator - 5 (Cat# R1013) kit. 1 µg of purified total RNA for each biological replicate was submitted to the Donnelly Sequencing Centre, upon which Poly-A-selected, rRNA-depleted libraries were generated with the TruSeq Illumina library preparation kit (Epicenter) according to the manufacturer’s instructions. The quantified library pool was hybridized to the flow cell to a final concentration of 2.21 pM and sequenced on the Illumina NovaSeq platform for 300 cycles (paired-end sequencing) to a target depth of 50 million reads.

### AlphaFold 2.0

AlphaFold 2.0 was used to model each protein, and AlphaFold-multimer was used to predict the structural contexts of the interactions between protein-protein pairs (Evans et al. 2022; Jumper et al. 2021). Full-length ZN121_HUMAN (P58317, https://www.uniprot.org/uniprotkb/P58317/entry) and YTHD2_HUMAN (Q9Y5A9, https://www.uniprot.org/uniprotkb/Q9Y5A9/entry) protein sequences were used for AlphaFold input.

AlphaFold 2.0 model parameters under Creative Commons Attribution 4.0 license were used for modelling of both individual proteins and co-structures. mmseq2 (Steinegger and Söding 2017) was used for Multiple Sequence Alignment (MSA) prior to prediction, and side-chain bond geometry was refined with Amber (Case et al. 2023). The highest-confidence models of individual proteins and pairwise co-structures (Case et al. 2023) were selected by their pLDDT and pTMscores, respectively, as calculated with AlphaFold. All visualizations were generated with Pymol and Chimera (Meng et al. 2023) (The PyMOL Molecular Graphics System, V1.2r3pre, Schrödinger).

### CLIP Methods

#### CLIP

All CLIP-based methods follow similar steps as described in (Nabeel-Shah, Pu, Burke, et al., 2024; Nabeel-Shah, Pu, Burns, et al., 2024; Nabeel-Shah & Greenblatt, 2023). HEK293 cells were grown to confluence in separate 15 cm plates for each biological replicate. After a 24h incubation with doxycycline (1μg/ml), cells were crosslinked with 0.15 J/cm^2^ at 254 nm in a Stratalinker 1800. After cell lysis in 2.2 ml of iCLIP lysis buffer (50 mM Tris pH 7.5, 100 mM NaCl, 1% NP40, 0.1 SDS, and 0.5% sodium deoxycholate, supplemented with protease inhibitor from Roche, Cat# 0589279100), 1 ml of the lysate was incubated with Turbo DNase (Life Technologies, Cat # AM2238) and RNase I (Ambion, Cat # AM2294) at 37°C for 5 min to digest genomic DNA and fragment RNA to ∼ 250 nt. The optimal concentration of RNase I was experimentally determined prior to performing CLIP. For each 1 ml of lysate, 50 μl of Dynabeads Protein G magnetic beads (Invitrogen™, Cat # 10003D) was used for pre-clarification, rotating at 4°C for 2h prior to IP. 6 μg of the appropriate antibody and 60 μl of Dynabeads Protein G magnetic beads were incubated with the clarified lysate for 2-4 hours (or overnight).

#### iCLIPseq

2% of the input material (from one iCLIP replicate for a given target) was saved to generate size-matched input control libraries (SMI) before IP. For GFP-ZNF121, a 1:250 RNase I dilution was used. Following the IP, beads were washed 5 times with iCLIP high salt buffer (50 mM Tris pH 7.5, 1 M NaCl, 1 mM NP40, 0.1% SDS, and 0.5 % sodium deoxycholate) with end-to-end rotation at 4°C, 5 min each, followed by three brief washes with PNK buffer (20 mM Tris pH 7.5, 10 mM MgCl2, 0.2% Tween-20). RNA dephosphorylation with FastAP was performed for 15 min at 37°C while shaking, followed by T4 polynucleotide kinase treatment for an additional 20 minutes, as described in the eCLIP procedure (Van Nostrand et al., 2016). Beads were washed 3 times with iCLIP high salt buffer for 5 min with end-to-end rotation at 4°C. Pre-adenylated iCLIP adaptors were ligated to the 3’-ends of RNA via the eCLIP ligation method (Van Nostrand et al. 2016). Beads were again washed with iCLIP high salt buffer three times for 5 min with end-to-end rotation at 4°C. Beads were subsequently washed three times with PNK buffer, and the IP’d RNA was 5’-end-labled with ^32^P with T4 polynucleotide kinase (New England Biolabs®, Cat# M0201L), separated using 4–12% NuPAGE BisTris–PAGE (Invitrogen™, Cat# NP0321BOX) and transferred overnight to a nitrocellulose membrane (Pall Corporation, Cat# 66485) in a cold room.

The inputs were also resolved on 4–12% BisTris–PAGE and transferred to a nitrocellulose membrane along with the IP’d RNA. The membrane was cut to match the size of the IP’d material, and the RNA was liberated from the membrane with Proteinase K (Thermo Fisher, Cat# 25530049) digestion. Reverse transcription was performed using barcoded primers, the cDNA was size-selected (low: 70 to 85 nt, high: 110 to 180 nt) using gel extraction (6% TBE-urea gels, Invitrogen, CAT # EC62652BOX), and the cDNA was circularized with CircLigase^TM^ II ssDNA ligase to ligate the adaptor to the 5’ end. Betaine was added to the CircLigase^TM^ reaction to 1 M final concentration, and the solution was incubated for 2h at 60°C. The circularized cDNA was linearized by cleavage at the internal BamHI site, followed by PCR amplification with AccuPrime SuperMix I (Thermo Fisher, Cat# 12342010). We performed agarose gel purification of the PCR products (Qiagen, Cat# 28106), and the eluted DNA was mixed at a ratio of 1:5:5 from the low, middle, and high fractions. This mixture was submitted for sequencing on either an Illumina HiSeq 2500 or Illumina NextSeq 500 platform using a High-Output v2.5 flow cell to generate single-end 51 nucleotide reads with 40M read depth per sample. The barcoded primers used for iCLIPseq are as follows: Rt9clip/5Phos/NNGCCANNN, sRt14clip/5Phos/NNTGCCNNN, Rt2clip/5Phos/NNACAANNNAGATCGGAAGAGCGTCGTGgatcCTGAACCGC, Rt15clip/5Phos/NNTATTNNNAGATCGGAAGAGCGTCGTGgatcCTGAACCGC, Rt16clip/5Phos/NNTTAANNNAGATCGGAAGAGCGTCGTGgatcCTGAACCGC.

Rt7clip/5Phos/NNCTAANNNAGATCGGAAGAGCGTCGTGgatcCTGAACCGC was utilized for SMIs.

#### CLIP RT-qPCR

Lysates were treated with 20 µl of RNase I (Ambion, Cat# AM2294) diluted to 1:500 (optimal for YTHDF2) and 2 µl of TURBO DNase (Life Technologies, Cat# AM2238). 10% of the clarified supernatant was kept as an input. 5 µg of anti-YTDHF2 antibody (Proteintech, Cat# 24744-1-AP) or rabbit-IgG as a control (Invitrogen™, Cat# 10500C) was incubated with the lysate overnight, rotating at 4°C, with 50 µl Dynabeads Protein G magnetic beads (Invitrogen™, Cat#10003D), pre-washed with CLIP lysis buffer added to the sample the next day for a 4°C, 2h incubation.

Samples were washed with high-salt CLIP wash buffer, rotated at 4°C, and briefly washed with PNK buffer and then PK buffer. Beads were resuspended in 100 µl of PK buffer (100 mM Tris pH 7.5, 50 mM NaCl, 10 mM EDTA) and treated with 50 µg of Proteinase K (Thermo Fisher Scientific, Cat# EO0491) at 55°C for 1 hour while shaking. The supernatant was supplemented with 300 µl of TE buffer (Invitrogen™, Cat# AM9849). RNA was extracted using one volume of Phenol:Chloroform (BioShop Canada Inc. Cat# PHE512.400) at room temperature for 10 min, then centrifuged at 15,000 x g for 10 min. To the aqueous phase was added 40 µl of 3 M sodium acetate (Invitrogen™, Cat# AM9740), 30 µg of GlycoBlue™ Coprecipitant (Thermo Fisher Scientific, Cat# AM9516), 1 ml of ice-cold anhydrous ethanol (Commercial Chemicals, Cat#P006EAAN), and incubated overnight at -20°C. Samples were centrifuged at 15,000x g for 15 min, and the pellets were washed with ice-cold 80% ethanol, and centrifuged at 15,000x g for 10 min. After allowing the pellets to air-dry for 3 min, the RNA pellets were rehydrated in DEPC-treated water. Input samples were processed using TRIzol Reagent (Invitrogen™, Cat# 15596026, 15596018) according to the manufacturer’s instructions.

For reverse transcription, 10 nM of dNTPs and random hexamers were incubated with the RNA sample at 70°C for 5 min. SuperScript III (Invitrogen™, Cat# 18080093) was utilized for reverse transcription. cDNA was subsequently diluted 1:10 for qPCR. Enriched RNA was analyzed relative to 10% of the input RNA material, and any IgG signal from the RT-qPCR was subtracted from the YTHDF2-IP to remove non-specific signal from the analysis. Primers designed for this experiment are listed in Supplemental Table 9.

#### CLIP-Autoradiography

Lysates were sonicated twice, 20 pulses per ml of lysate on ice (0.3 seconds on, 0.7 seconds off, 30% amplitude). 20µl of RNase I (Ambion, Cat# AM2294) 1:250 in CLIP lysis buffer, unless otherwise indicated, and 4 µl of TURBO DNase (Life Technologies, Cat# AM2238) were added and were shaken with samples for 5 min at 37°C. Following 3 min on ice, samples were clarified for 20 min at 16,000x g, 4°C and the IP was carried out as described above.

Samples were washed 6 times with 900 μl high-salt CLIP wash buffer and incubated on a rotator at 4°C for 5 min. Samples were then washed briefly with 900 μl PNK buffer and then 900 μl PK buffer. Beads were resuspended in 100 µl of PK buffer.

IP’d RNA was 5’end-labeled using T4 polynucleotide kinase and ^32^P-ATP (New England Biolabs^®^, Cat# M0201L), then protein-RNA complexes were separated on a 4-12% BisTris-SDS-PAGE gel and transferred overnight to a nitrocellulose membrane (Pall Corporation, Cat# 66485) at 4°C. The membrane was then exposed to X-ray film for RNA detection.

#### Nuclease over-digestion assay

As in the CLIP-Autoradiography procedure. However, after IP and stringent washes using high salt wash buffer and RNA 5’-end-labeling with ^32^P using T4 polynucleotide kinase, equivalent volumes of beads were incubated with 5 μL of RNase I or Turbo DNase for 15 min while shaking (1200 rpm) at 37 °C. Beads were washed once with 900 μL of PNK buffer. Protein-RNA complexes were separated using 4–12% BisTris– SDS-PAGE and transferred to a nitrocellulose membrane (Pall Corporation, Cat# 66485), followed by autoradiography.

### iCLIPseq data analysis

iCLIPseq data was analyzed as described in (Nabeel-Shah et al. 2024b). Briefly, ≥51-nt iCLIPseq raw reads consisting of three random positions, a 4-nt multiplexing barcode, and another two random positions, followed by the cDNA sequence, were de-duplicated based on the first 45 nt. Reads were de-multiplexed, and we discarded random positions, barcodes, and any 30 nt matching Illumina adaptors, and reads shorter than 25 nt. The remaining reads were trimmed to 35 nt using Trimmomatic (Bolger et al. 2014) and mapped to the human genome/transcriptome (Ensembl annotation of hg19) with Tophat (Trapnell et al. 2009) on default settings. To prevent false read assignments from repetitive regions, reads with a mapping quality of < 3 were discarded.

miCLIPseq and YTHDF2 iCLIPseq raw data were obtained from (Nabeel-Shah et al. 2024a, 2024b), and can be obtained via the NCBI Gene Expression Omnibus (https://www.ncbi.nlm.nih.gov/geo/) with the accession number GSE228608 and GSE230846.

### Identification of crosslink-induced truncation sites (CITS)

We defined RNA-binding sites at single nucleotide resolution (Nabeel-Shah et al. 2024b). We used the CLIP Tool Kit (CTK) method (Shah et al. 2017) and called CITS on CLIP replicates and individual SMI samples. We only considered CITS with *P* < 0.01 for downstream analysis. Further, we removed our previously identified crosslinking hotspots (Nabeel-Shah, Pu, Burns, et al., 2024) for downstream analysis. The remaining CITS were normalized using SMI CITS, and underlying reads from the iCLIP CITS and SMI CITS were used to perform differential analysis using the R package DEseq2, with a cut off *P* < 0.05. miCLIPseq data was similarly processed, and putative m6A sites within the DRACH motif were considered miCLIPseq CITS. We plotted the CITS along mRNA transcripts by generating a metagene plot with the R package GenomicPlot (URL: https://github.com/shuye2009/GenomicPlot), and MetaPlotR (Olarerin-George and Jaffrey 2017). Peak distributions across genomic regions (analyzing 5’ and 3’UTRs separately) were calculated using ‘GenomicPlot’.

### Overlap of ChIPseq and iCLIPseq reads

To compare the RNA- and DNA-binding positions of ZNF121, we correlated the ZNF121 iCLIPseq reads with previously published ZNF121 ChIPseq data (Schmitges et al., 2016), GEO GSE76496. ZNF121 iCLIPseq read density around ZNF121 ChIPseq peaks was computed and plotted using the R package ‘GenomicPlot’.

### Comparison of miCLIPseq for m6A sites with GLORI for m6A sites

miCLIPseq m6A sites were overlapped with GLORI m6A sites using the ‘intersect_bed’ function in the R package ‘valr’ <doi:10.12688/f1000research.11997.1>. The methylation levels of the overlapping sites were compared to ‘clustered’ and ‘non-clustered’ GLORI m6A sites as a density plot using the ‘geom_density’ function in the R package ‘ggplot2’.

### Comparison of ZNF121 and YTHDF2 iCLIPseq sites with GLORI m6A sites

ZNF121 and YTHDF2 iCLIPseq CITS, as well as GLORI m6A sites, were resized to 100 nt, and then the overlaps of ZNF121 and YTHDF2 iCLIPseq CITS with GLORI m6A sites were calculated and displayed as Venn diagrams using the R package ‘GenomicPlot’. GLORI data was downloaded from the NCBI Gene Expression Omnibus (GEO) using the GEO accession number <u>GSE210563</u>.

### RNAseq analysis

To determine differential expression, reads were aligned to the human genome GRCh38 using STAR (version 2.7.6.a)(Dobin et al. 2013), gene level read counts were quantified by RSEM (version 1.3.3), and DESeq2(Love et al. 2014) was used to characterize the data by variance-stabilizing transformation. Adjusted p-value ≤ 0.05 and absolute log2fold change ± 1 were used to identify significantly regulated genes.

### icSHAPE, RNAplFold, and RNA structural prediction using in silico folding of ZNF121 target sequences

iCLIPseq-determined targets of ZNF121 were overlapped with transcriptome-wide icSHAPE nucleotide reactivity scores in HEK293 cells and RNAplFold in silico folding predictions using the pipeline described (Nabeel-Shah et al. 2024b).

icSHAPE data: (GEO: GSE117840)

Scripts for the icSHAPE/iCLIP overlap pipeline are publicly available through GitHub: https://github.com/jburnsuoft/icSHAPE_iCLIP.

https://zenodo.org/records/13375429

## Acknowledgements

We acknowledge Drs. Edyta Marcon, Zuyao Ni, and Ernest Radovani, as well as Hua Tang, for their insights, helpful discussion, and technical assistance. We also thank Drs. Ulrich Brauchweig and Benjamin J. Blencowe, as well as Hyunmin Lee and Dr. Zhaolei Zhang, for help with iCLIPseq data analysis. We also thank Sherin Shibin and the team at The Donnelly Sequencing Centre for their assistance with next-generation sequencing.

## Author Contributions

G.L.B and J.F.G conceived this study. G.L.B designed the experiments with input from S.N-S, S.M.M and J.F.G. G.L.B coordinated the project, wrote the manuscript with J.F.G, and performed all experiments (except as outlined below). S.N-S performed and helped analyze all CLIP-autoradiography and iCLIPseq experiments. S.P performed all computational bioinformatics analyses related to iCLIPseq, RNAseq, and ChIPseq experiments. N.A generated CRISPR-KO cell lines for ZNF121 and YTHDF2, and helped perform iCLIPseq and CLIP-autoradiography experiments. S.M.M helped with western blots and performed immunofluorescence experiments. J.B performed the AlphaFold 2.0 and icSHAP/RNAplFold analyses. A.A helped with cell growth assays, and A.A and G.Z helped with cell line construction. All authors participated in editing the manuscript and contributing to the Materials and Methods.

## Data Availability

Sequencing data for RNA-seq and iCLIP-seq experiments have been deposited in the European Nucleotide Archive (ENA) (https://www.ebi.ac.uk/ena/browser/home) under accession number PRJEB.

## Supplemental Figure Legends

**Supplemental Figure 1.** ZNF121 interacts with YTHDF2, and presence of ZNF121 and YTHDF2 in cytoplasmic and nuclear extracts. (*A*) Gene Ontology (GO) Biological Processes term enrichment for proteins which putatively interact with ZNF121 in our analyzed AP-MS data (BFDR ≤ 0.01). (*B*) Western blots showing co-IP of YTHDF2 from extracts of HEK293 cells expressing GFP fusion proteins after IP with anti-GFP antibodies of GFP-ZNF121. Cells expressing free GFP are negative controls. (*C*) Western blot showing co-IP of GFP-ZNF121 from extracts of HEK293 cells expressing free GFP or GFP-ZNF121 after IP with anti-YTHDF2 antibodies. (*D*) Western blot showing co-IP of YTHDF2 after IP with anti-GFP from extracts of HEK293 cells expressing GFP or GFP-ZNF121 and treated with the indicted amounts of RNase I. (*E*) Western blot of nuclear and cytoplasmic fractions from GFP-ZNF121-expressing HEK293 cells, probing for GFP, YTHDF2 and the nuclear and cytoplasmic markers, DEK and GAPDH, respectively. (*F*) As described in *E*, utilizing nuclear and cytoplasmic fractions from normal HEK293 cells and probing for endogenous ZNF121.

**Supplemental Figure 2**. The C-terminus of YTHDF2 facilitates an interaction with the ZnFs of ZNF121. (*A*) Ribbon diagram of the structure of the ZNF121-YTHDF2 heterodimer predicted by AlphaFold multimer, using the structures of ZNF121 (pink) and YTDHF2 (purple) predicted by AlphaFold 2.0. Only the interacting globular domains are depicted for clarity. Numbers refer to the ZnFs of ZNF121. (*B*) Predicted local distance difference test (pLDDT) plot, depicting the local confidence of each residue in the predicted AlphaFold 2.0 structures from the top five ranking structures generated for each protein. Positions 0-579 are the pLDDT for YTHDF2 (note that residues 0-399 are considered disordered) and positions 580-970 are the pLDDT for ZNF121 (note that residues 580-668 are considered disordered). (*C*) Predicted aligned error (PAE) plot for YTHDF2 and ZNF121. PAE measures the confidence of AlphaFold 2.0 in the positioning of any two residues relative to each other in the predicted structure, measured as an error rate for each aligned residue. Red indicates an increased error rate (less confidence) in the positioning of two given residues, whereas blue (higher confidence) represents a lower error rate in the prediction. Chain A is YTHDF2, and chain B is ZNF121. Aligned residues from chains A or B are compared to scores from the YTHDF2 peptide chain (0-579) and the ZNF121 peptide chain (580-970) (Y-axis). (*D*) Rendering of the predicted ZNF121-YTHDF2 predicted complex, disordered regions included, by chain (left; blue represents ZNF121 and green represents YTHDF2) as a reference, or by pLDDT, with red signifying less confidence and blue signifying greater confidence. (*E*) Alignment of our predicted structure of the ZNF121 (pink) and YTHDF2 (purple) complex with a structure of YTHDF2 complexed with a m6A-C-U RNA sequence resolved by X-ray diffraction (lavender and green, respectively). The m6A binding pocket residues and RNA interacting residues of the predicted YTDHF2 structure are highlighted in yellow and red, respectively. Numbers refer to the ZnFs of ZNF121. (*F*) Layout of the YTH-domain of the YTHDF paralogues relative to YTHDF2. Arrows represent β-sheets, and rectangles represent α-helices. Residues highlighted in red represent those which are conserved among all three YTHDF paralogues. Purple stars represent those residues that form the tryptophan cage. Blue stars indicate residues important for RNA binding. (*G, Top*) ZNF121, and (*Bottom*) YTHDF2 protein organization, showing the various truncations used in various experiments. (*H,I*) Western blots showing co-IP of YTHDF2 after IP with anti-FLAG antibodies from extracts of HEK293 cells expressing the indicated FLAG-ZNF121 constructs, where FL is the full-length ZNF121, NT is residues 1-171, M is 167-283, CT is 279-390, and ΔIDR is 88-390. (*J*) Western blot showing co-IP of ZNF121 after IP with anti-GFP antibodies of GFP-YTHDF2 fusion proteins from extracts of HEK293 cells expressing the indicated YTHDF2 fusion constructs. (*K*) Western blot showing co-IP of ZNF121 from cell extracts after IP with anti-GFP antibodies of FLAG-YTHDF1, FLAG-YTHDF2, or FLAG-YTHDF3 fusion proteins.

**Supplemental Figure 3**. iCLIPseq identifies ZNF121 as an RBP that primarily binds 3’UTRs near YTHDF2 binding sites. (*A*) SDS-PAGE and subsequent silver staining of proteins IP’d under CLIP or standard co-IP conditions with anti-GFP antibodies from lysates of HEK293 cells expressing GFP-ZNF121. (*B*) Over-digestion assay. (*Left*) schematic of the over-digestion assay. (*Right*) Autoradiograph of ^32^P-labeled RNA derived from UV-crosslinked HEK293 cells expressing GFP-ZNF121 following over-digestion of the IP’d material with RNase I or Turbo DNase, as indicated. Below is a corresponding Western blot probing for GFP-ZNF121. (*C*) Schematic of GFP-ZNF121 constructs used in CLIP-autoradiography experiments. (*D*) Assignment of GFP-ZNF121 iCLIPseq CITS to the indicated RNA species as a percentage of the total. (*E*) Red: distribution of GFP-ZNF121 iCLIPseq ratio-over-input around the summit ChIPseq peaks of GFP-ZNF121. Blue depicts random sites within 200 nt of the ZNF121 iCLIPseq peak. (*F*) Overlap of genes bound by ZNF121 at the transcript level, based on iCLIPseq data, or their promoter DNA, based on ChIPseq data. *p* was calculated using the Hypergeometric Distribution. (*G*) Top five RNA motifs identified in sequences bound by ZNF121 utilizing the DREME suite (Bailey, 2011). (*H*) Prediction of RNA structure at centers of ZNF121 iCLIPseq crosslinking sites (vertical dashed line). *In vivo* (+ protein, orange) and *in vitro* (– protein, blue) are icSHAPE-generated (Flynn et al. 2016) nucleotide reactivity scores. Solid grey represents the RNAplFold-predicted (Lorenz et al. 2011) average unpaired probability score, functioning as an *in silico* analogue to icSHAPE. (*I*) Venn diagram showing the overlap of YTHDF2 CITS with ZNF121 CITS located within 100 nt of each other in the CDS. Jaccard similarity coefficient was used to determine the degree of similarity between the two data sets.

**Supplemental Figure 4**. Knockout of ZNF121 results in changes to mRNA abundance of bound targets. (*A*) Western blot validating the CRISPR/Cas9 KO of ZNF121 relative to a GAPDH loading control (upper panel), and the associated quantification (lower panel). (*B*) Volcano plot for differential mRNA abundance upon ZNF121 KO relative to a wild-type control. Each dot represents a transcript. Log2Fold cut off of ±1 and a *padj* of ≤ 0.05 were utilized to identify significantly differentially expressed transcripts. (*C*) Gene-set enrichment analysis of differentially expressed transcripts upon ZNF121 KO. (*D*) Proportion of up- or down- regulated transcripts from ZNF121 KO RNAseq that were bound by ZNF121 at the RNA level (based on iCLIPseq data) or on promoter DNA (based on ChIPseq data), or both, and whether they are up-or down-regulated upon ZNF121 KO. Targets with Log2Fold expression change cut off ± 0.05 and *Padj* of ≤ 0.05 were utilized. (*E, Left*) Western blots of ZNF121 and YTHDF2 in WT-HEK293 cells or two ZNF121 KO clones (clone 1 represents a partial KO and clone 2 represents a full KO used in all subsequent experiments). (*Right*) The corresponding quantification for YTHDF2 relative to GAPDH. Data presented as ±SEM, n=3. *p,* ≤ 0.05 (*); ≤ 0.005 (**); ≤ 0.001 (***); > 0.05 (n.s) using Student t-test.

**Supplemental Figure 5**. The mRNA binding sites of ZNF121 mostly exclude m6A sites, and most mRNA binding sites of ZNF121 and YTHDF2 do not have nearby m6A. (*A*) Distribution of m6A sites determined by miCLIPseq around YTHDF2 or ZNF121 CITS, with or without the other protein located within 100 nt, as indicated. (*B*) Distribution of ZNF121 CITS around nearby YTHDF2 CITS in transcripts with or without m6A within 100 nt, as determined by miCLIPseq, across all transcript features. (*C*) Numbers of transcripts bound by ZNF121 and YTHDF2 with CITS located within 100 nt of each other in all exons or only in 3’UTRs. (*D*) Overlap of m6A sites identified by our miCLIPseq data (Nabeel-Shah et al. 2024b, 2024a) with m6A sites identified by GLORI (Liu et al. 2022). Jaccard similarity coefficient was used to determine the degree of similarity between the two data sets. (*E*) Comparison of the methylation frequencies of m6A sites identified by GLORI with those identified by miCLIPseq data. When GLORI m6A sites occur near one another, they are represented as a single clustered GLORI m6A site, as opposed to non-clustered sites, in which each identified site is considered an individual m6A site. (*F-I*) Venn diagrams showing overlaps of ZNF121 or YTHDF2 CITS with m6A sites identified by miCLIPseq (*F-G*) or GLORI (*H-I*). (*J*) Venn diagram showing overlaps in transcripts of ZNF121 and YTHDF2 CITS and m6A sites identified by GLORI within 100 nt. *p* determined by the Hypergeometric Distribution. (*K-L*) Methylation frequencies of m6A sites identified by GLORI near YTHDF2 (*K*) or ZNF121 (*L*) CITS.

**Supplemental Figure 6**. ZNF121 affects mRNA stability by recruiting YTHDF2 to common transcripts, modulating target poly(A) tail length, and interacting with the PAN2-PAN3 complex. (*A*) YTHDF2 RIP-qPCR in ZNF121 KO or wild-type HEK293 cells, as indicated, for transcript targets containing YTHDF2 and ZNF121 binding sites, with (*left*) or without (*right*) m6A sites identified by miCLIPseq within 100 nt. Data presented as ±SEM, n=3. *p,* ≤ 0.05 (*); ≤ 0.005 (**); ≤ 0.001 (***); > 0.05 (n.s) using Student t-test. (*B*) Effects of ZNF121 KO on Poly(A)-tail lengths for *CASP8, MDM2,* and *CBLLI* mRNAs using the Affymetrix Poly(A)-tail length assay. (*C*) Western blot showing co-IP of YTHDF2, PAN2, and CNOT1 after IP with anti-GFP from extracts of HEK293 cells expressing GFP or GFP-ZNF121. (*D*) Western blot showing co-IP of CNOT1, PAN2, or ZNF121 after IP with anti-FLAG antibodies or IgG using extracts from HEK293 cells expressing FLAG-tagged YTHDF2.

**Supplemental Figure 7**. Knockout of ZNF121 results in increased MDM2 protein abundance and delayed DNA damage response. (*A*) KEGG enrichment analysis for those genes whose transcripts contain CITS for both ZNF121 and YTHDF2 in their 3’UTRs (*p < 0.05*). (*B*) STRING analysis of KEGG enriched p53-related pathways identified in *A*, including p53 signalling, cell cycle regulation, and cellular senescence. Edges represent physical interactions or genes involved in similar pathways. Edge thickness indicates the strength of the association between two nodes. (*C*-*D*) Effect of single or pooled siRNAs targeting CNOT1 (siCNOT1) (*C*) or PAN2 (siPAN2) (*D*) on their expression levels determined by RT-qPCR. (*E*, *Left*) Western blot showing the effect of the CRISPR/Cas9 knockout of YTHDF2 in HEK293 cells on the expression of YTHDF2 and ZNF121. (*Right*) Quantification of the effect on expression of YTHDF2. (F) Effect on *MDM2* transcript abundance, as determined by RT-qPCR and normalized to levels in untreated HEK293 cells, of knocking out YTHDF2 or treating the cells with siRNA against PAN2 or CNOT1. (*G*) Western blot showing the effects of knocking out ZNF121 in HEK293 cells on the abundance of MDM2, ZNF121, and p53. (*H*) Effect of knocking out ZNF121 in HEK293 cells on the relative transcript levels of p53 targets *p21* and *p27*. (*I, top*) Western blot showing H2A.X activation in the presence or absence of ZNF121 following treatment with 1 mM Hydroxyurea (HU) for 24h. (*bottom*) Corresponding quantifications. Data presented as ±SEM, n=3. *p,* ≤ 0.05 (*); ≤ 0.005 (**); ≤ 0.001 (***); > 0.05 (n.s) using Student t-test.

**Supplemental Figure 8.** Knockout of ZNF121 reduces cell growth, and ZNF121 expression is increased in colorectal adenocarcinoma. (*A*) Western blot showing H2A.X activation over time following treatment with 1 mM hydroxyurea (HU). (*B*) Quantification of western blot signal after probing with anti-γH2AX antibodies in WT-HEK293 or ZNF121-KO cells at the indicated times following HU treatment. Data presented as ±SEM, n=4. *p,* ≤ 0.05 (*); ≤ 0.005 (**); ≤ 0.001 (***); > 0.05 (n.s) using Student t-test. (*C*) Live-cell images showing growth over 96h of wild-type HEK293 cells or cells with ZNF121 knocked out. All wells were initially seeded with 6×10^3^ cells. (*D*) TCAG data from Gepia 2.0 (Tang et al. 2019) showing ZNF121 expression in colorectal adenocarcinomas (T) and their various subtypes (MSI-H – microsatellite instability high, MSI-L – microsatellite instability low, MSS – microsatellite stable) compared to ZNF121 levels in normal tissues (N). (*E*) Kaplan–Meier plots using TCAG data from Gepia 2.0 (Tang et al. 2019) showing survival of colorectal adenocarcinoma patients with high or low ZNF121 expression. Logrank refers to the logrank test. HR refers to the hazard ratio test. The number of patients included in these analyses are represented by n.

**Supplemental Figure 9.** Preliminary Western blots showing co-IP of YTHDF2 from extracts of HEK293 cells expressing GFP fusion proteins after IP with anti-GFP antibodies of GFP-ZNF224. Cells expressing free GFP are negative controls.

**Supplemental Table 1.** SAINTexpress scored putative ZNF121 interacting proteins by APMS: SAINTexpress results for ZNF121 AP-MS from HEK293 cells. Preys with BFDR ≤ 0.01 are considered high-confidence and highlight in Green. Bait protein is GFP tagged. Prey is the Gene Accession in NCBI Entrez Gene and PreyGene is the Official Gene Symbol. Spectral counts for the prey (column D separated by “I” delimiter; column E, summed, column F, averaged), number of replicates performed (column G), spectral counts for the prey across all negative controls (column H), averaged probability across replicates (column I), maximal probability (column J),SaintScore (column K), Fold Change (counts in the purification divided by counts in the controls plus small factor to prevent division by 0; column L) and Bayesian FDR (column M) are listed are directly from the SAINTexpress output. Samples were acquired on an Orbitrap mass spectrometer. Note: Bait spectral counts are omitted from the list.

**Supplemental Table 2.** Transcripts bound by ZNF121 based on Crosslinking induced trucation sites (CITS) from GFP-ZNF121 iCLIPseq: Only CITS that passed a threshold of FDR ≤ 0.01 were considered for downstream analysis. Column A reports the Ensembl transcript identification number (ID). Columns B-H show the number of ZNF121 CITS in each of the indicated gene features for each transcript ID. Column I indicates the total amount of ZNF121 CITS per transcript ID. Column J-L indicate the genomic coordinates of the ZNF121 CITS, from chromosome (J) and the start and end position (K,L). Column M represents the strand for which the CITS were identified. Column N lists the Ensembl gene ID’s and Column O lists the gene names. Lastly, Column P lists the RNA biotype of the transcripts for which ZNF121 CITS were identified.

**Supplemental Table 3**. Transcript targets containing ZNF121 and YTHDF2 CITS (BFDR ≤0.01). Listed in column A are the transcripts that are bound by both ZNF121 and YTHDF2 in exons. Column B lists those transcripts that contain ZNF121 CITS in exons, but not YTHDF2 CITS. Column C list those genes that contained CITS for YTHDF2, but not ZNF121 in exons. Columns D-E are the same as A-C, however, consider CITS exclusively in 3’UTRs. This table supplements Figure 3E.

**Supplemental Table 4.** Differential expression of genes in ZNF121-KO relative to WT-HEK293 cells. DEseq2 analysis of RNAseq using RNA extracted from ZNF121 KO relative to that of WT-HEK293 cells. Column A indicates the Ensembl gene ID, and colum B shows the corresponding gene name. The base mean is listed in column C, and Log2Fold change is calculated in column D. The Log2Fold change standard error (SE) is indicated in column E. The p value is calculated in column G and the adjusted p value, taking into account multiple hypothesis testing is listed in column H. RNAseq was performed on the Illumina NovaSeq 6000.

**Supplemental Table 4**. Log2Fold change in expression of m6A-related targets upon ZNF121 KO, determined by RNAseq. Log2Fold change ± 0.05 and *Padj* of ≤ 0.05 were utilized to determine a significant change.

**Supplemental Table 5**. RNAseq differential expression of m6A-related transcript in the presence or absence of ZNF121. Colum A indicates the role each gene plays in m6A biology. Column B lists each m6A related gene. Column C indicates the Log2 fold change, and column D identifies the adjusted p value, correcting for multiple hypothesis. The significance differential expresion is expressed in column E as Log2Fold change cutoff ≥ 0.05 and ≤ - 0.05; * *Padj* of ≤ 0.05, ** *Padj* ≤ 0.005, *** *Padj* ≤ 0.001.

**Supplementary Table 6.** Transcript targets containing ZNF121 and YTHDF2 CITS: Counts of m6A sites identified by miCLIPseq, as well as ZNF121 CITS and YTHDF2 CITS, either in all exons or only in the 3’UTRs of transcript targets bound by either ZNF121 and YTDHF2, or only by YTHDF2. The intensity of the red indicates the relative amount of CITS/m6a sites.

**Supplemental Table 7**. Primers used for plasmid construction and cloning experiments.

**Supplemental Table 8**. Antibodies utilized in IP, Western Blot, RIP, CLIP or immunofluorescence experiments.

**Supplemental Table 9**. Primers used for RT-PCR and RT-qPCR experiments.

