## Supplemental Figures 1-8 for "ZNF121 recruits YTHDF2 to modulate mRNA stability"

### Supplemental Fig. S1

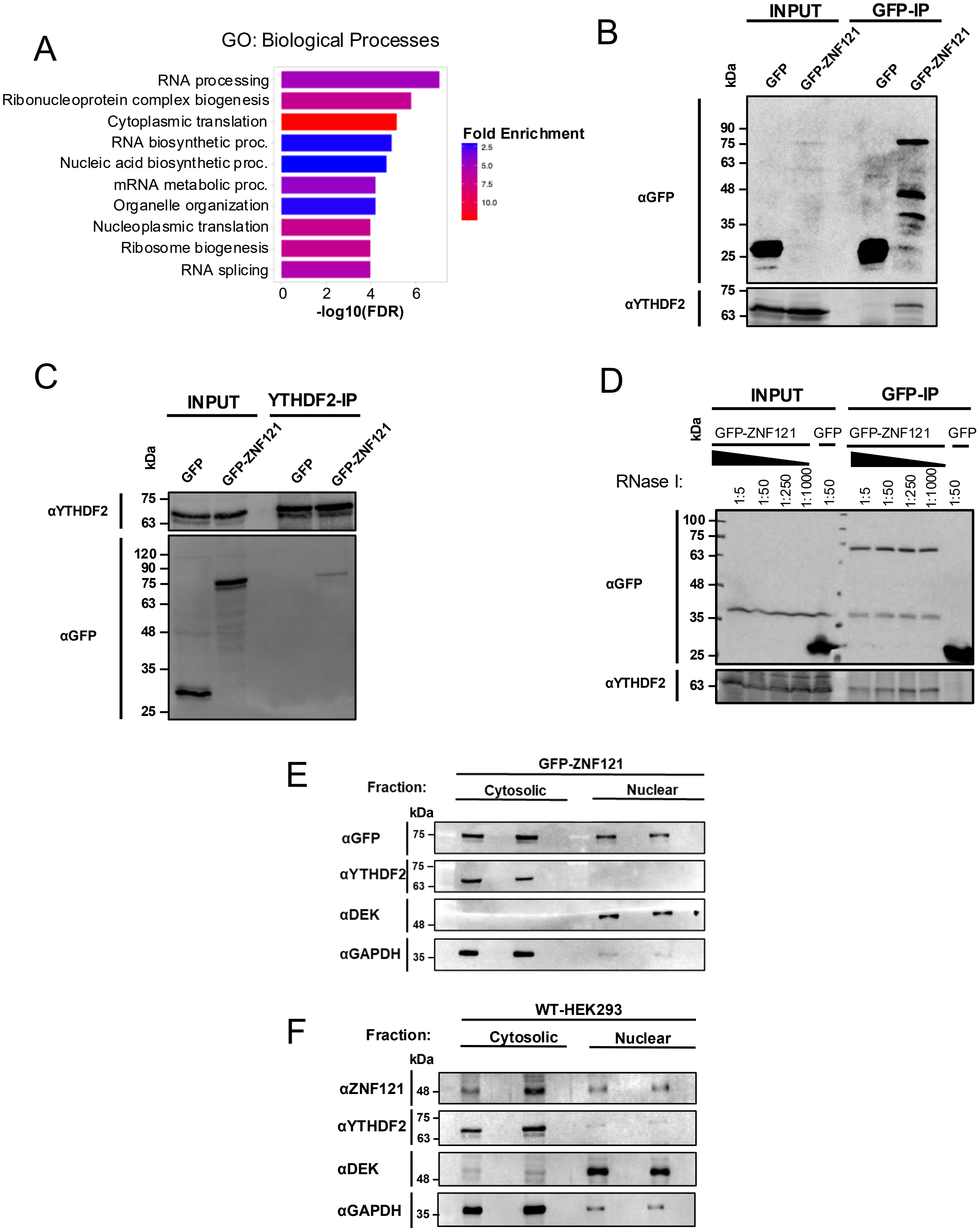

### Supplemental Fig. S2

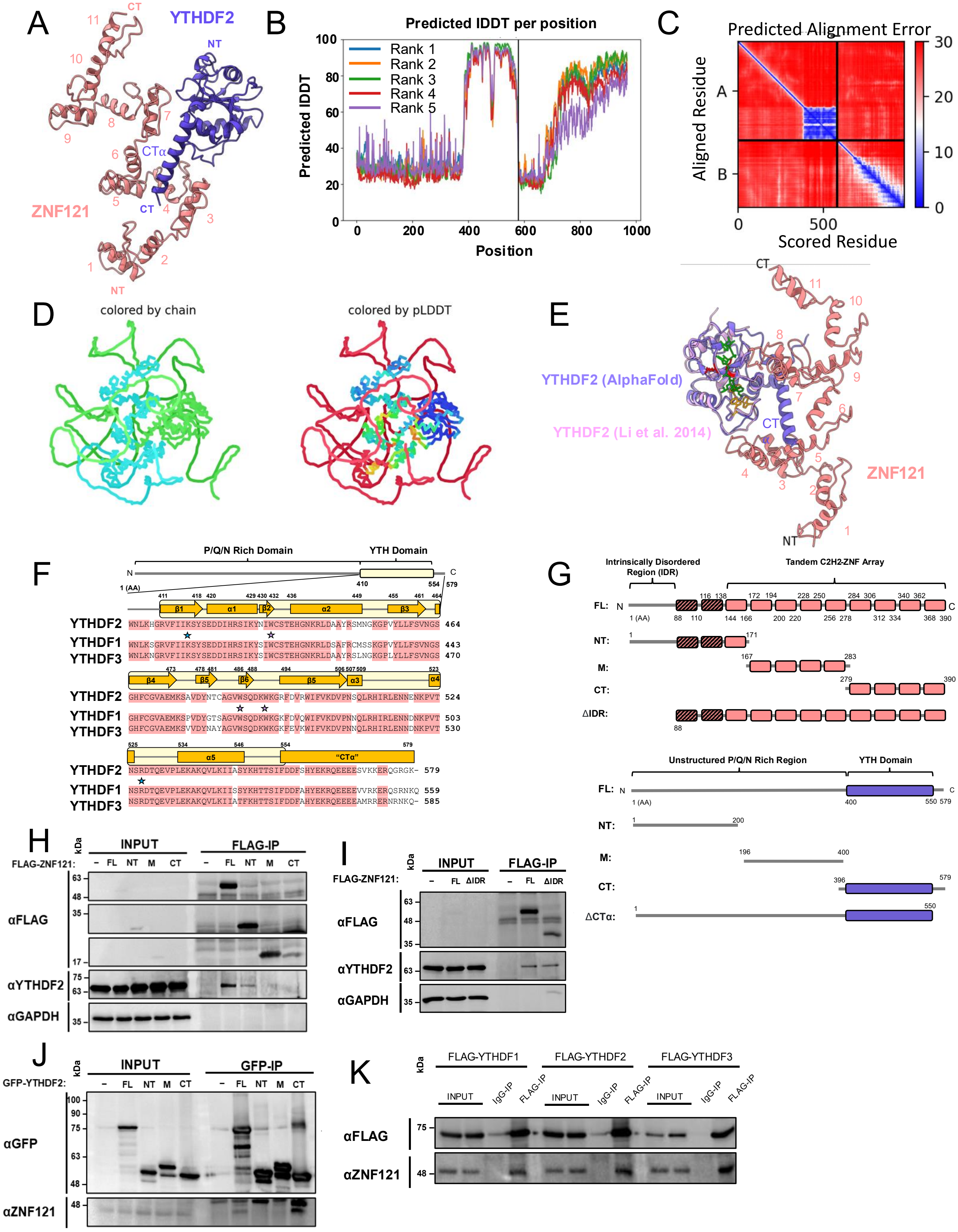

### Supplemental Fig. 3

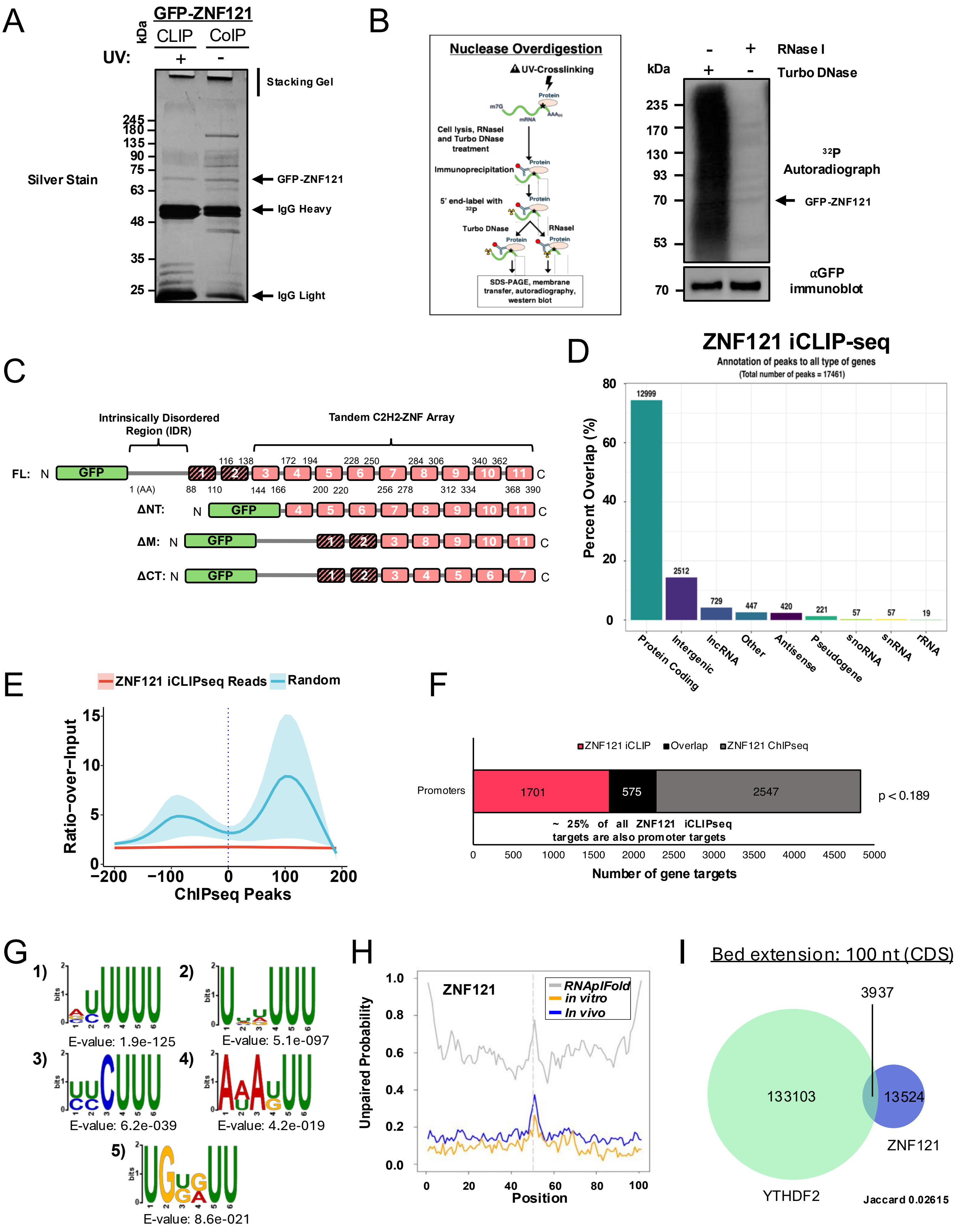

### Supplemental Fig. S4

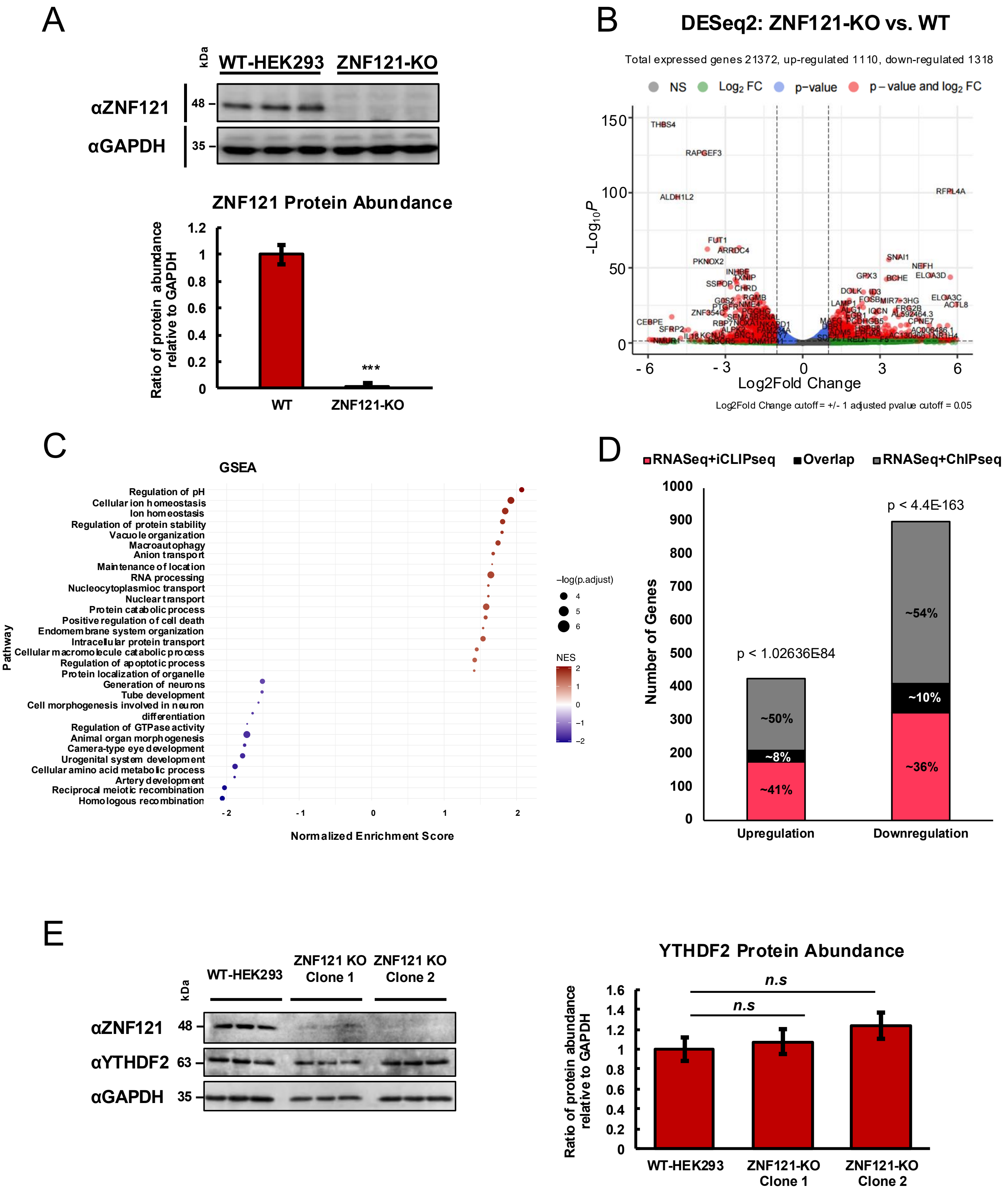

### Supplemental Fig. 5

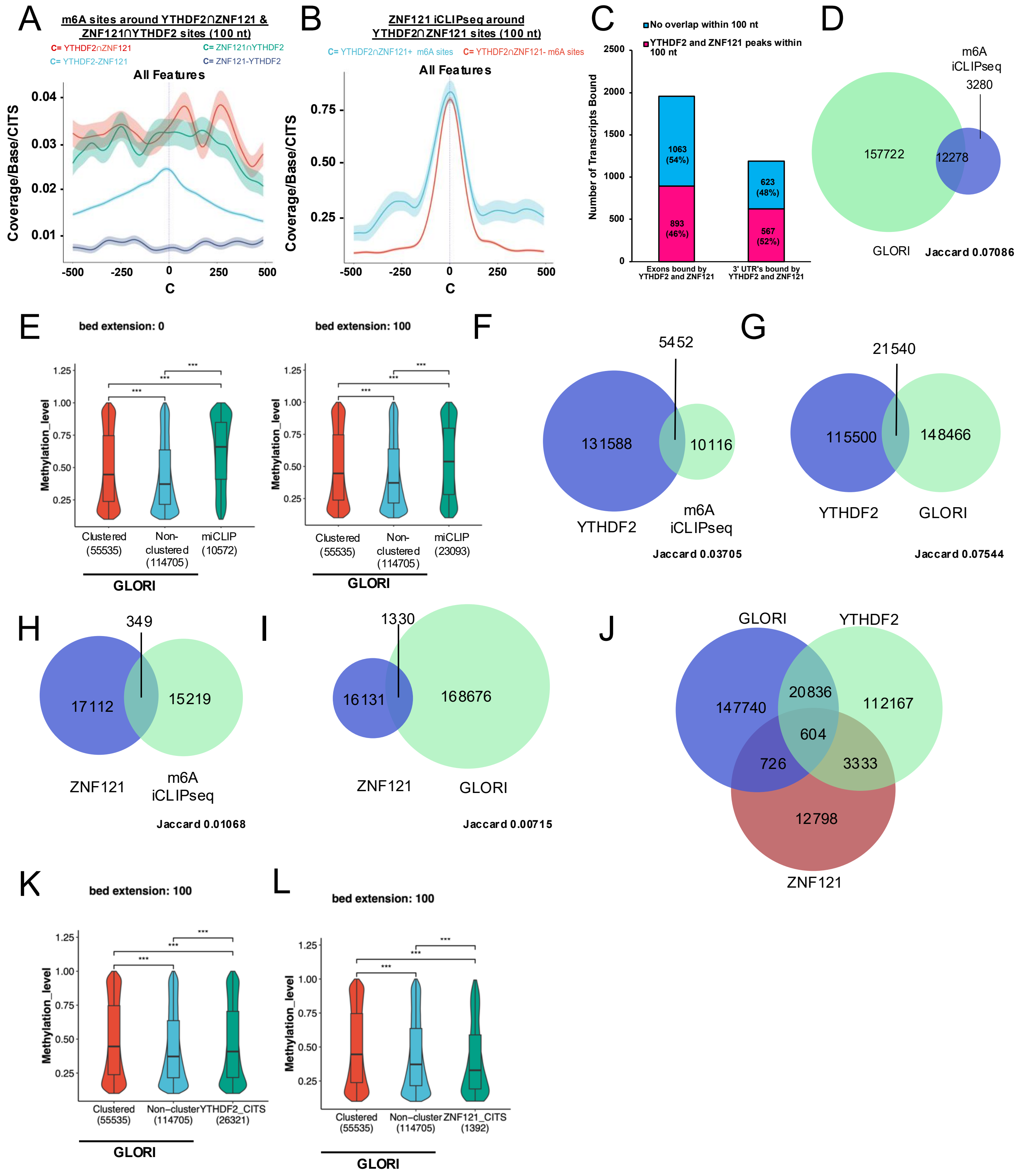

### Supplemental Fig. 6

A

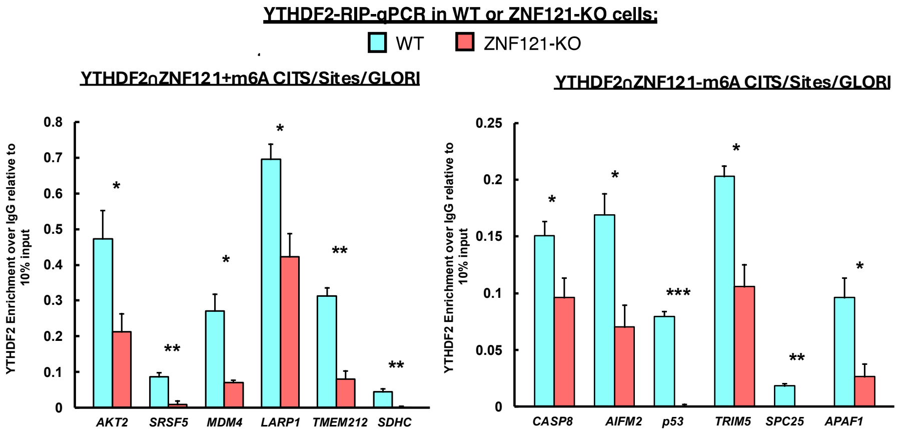


B

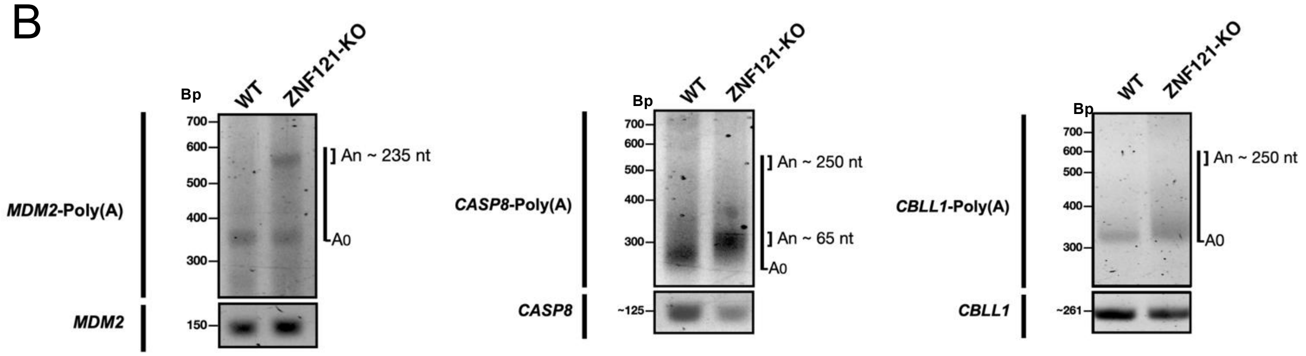


C

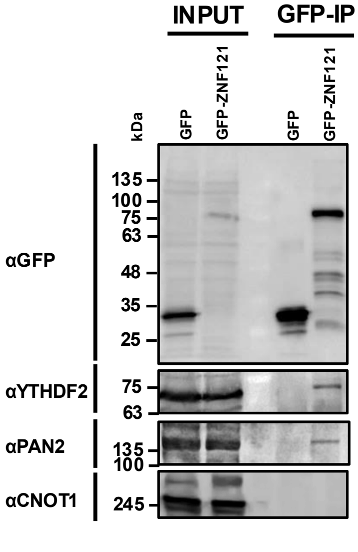


D

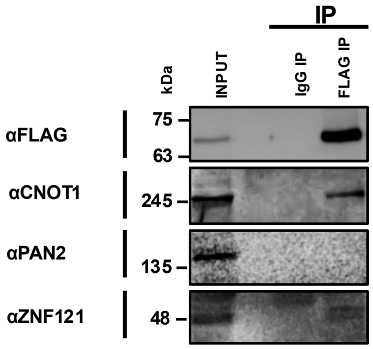

### Supplemental Fig. 7

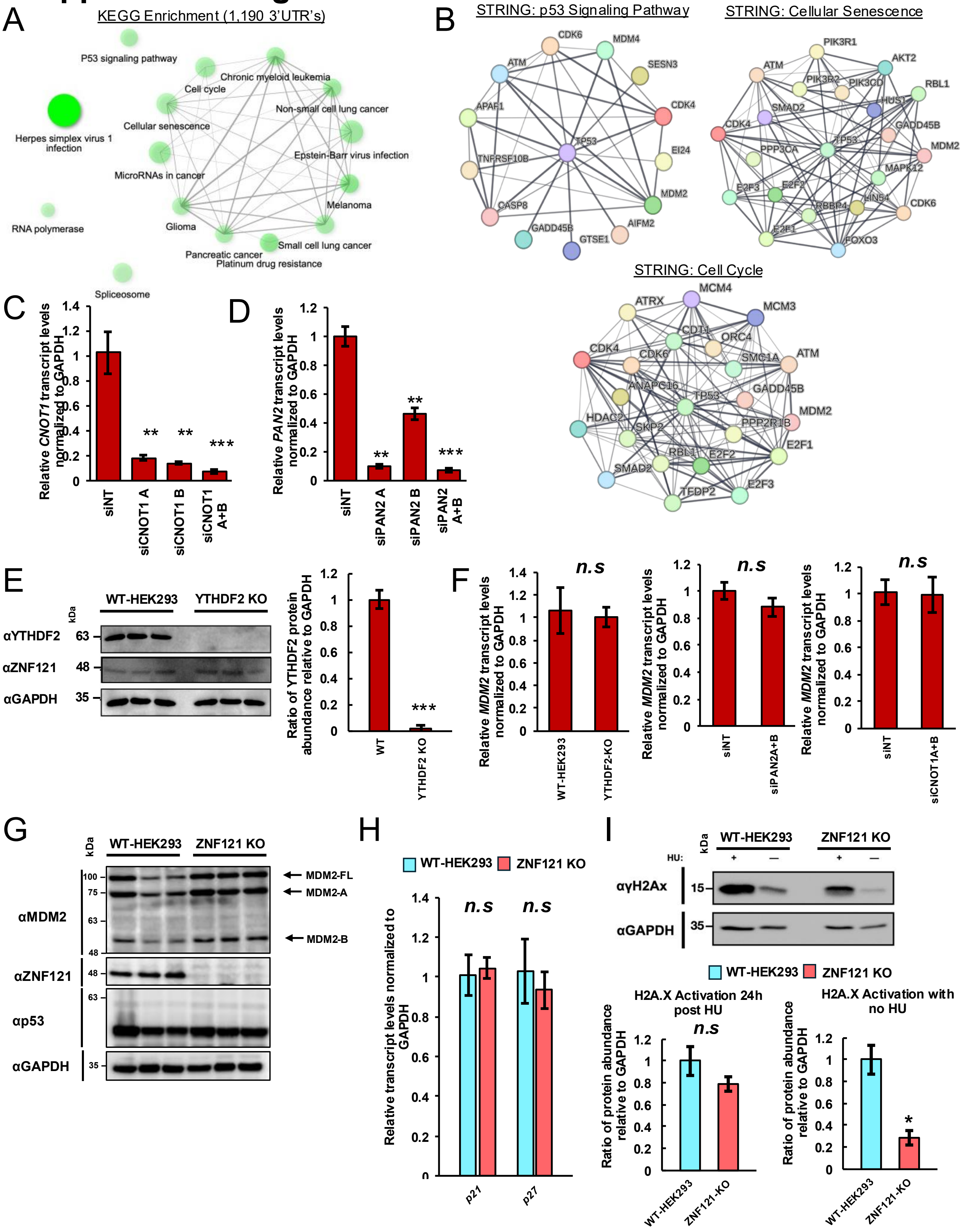

### Supplemental Fig. 8

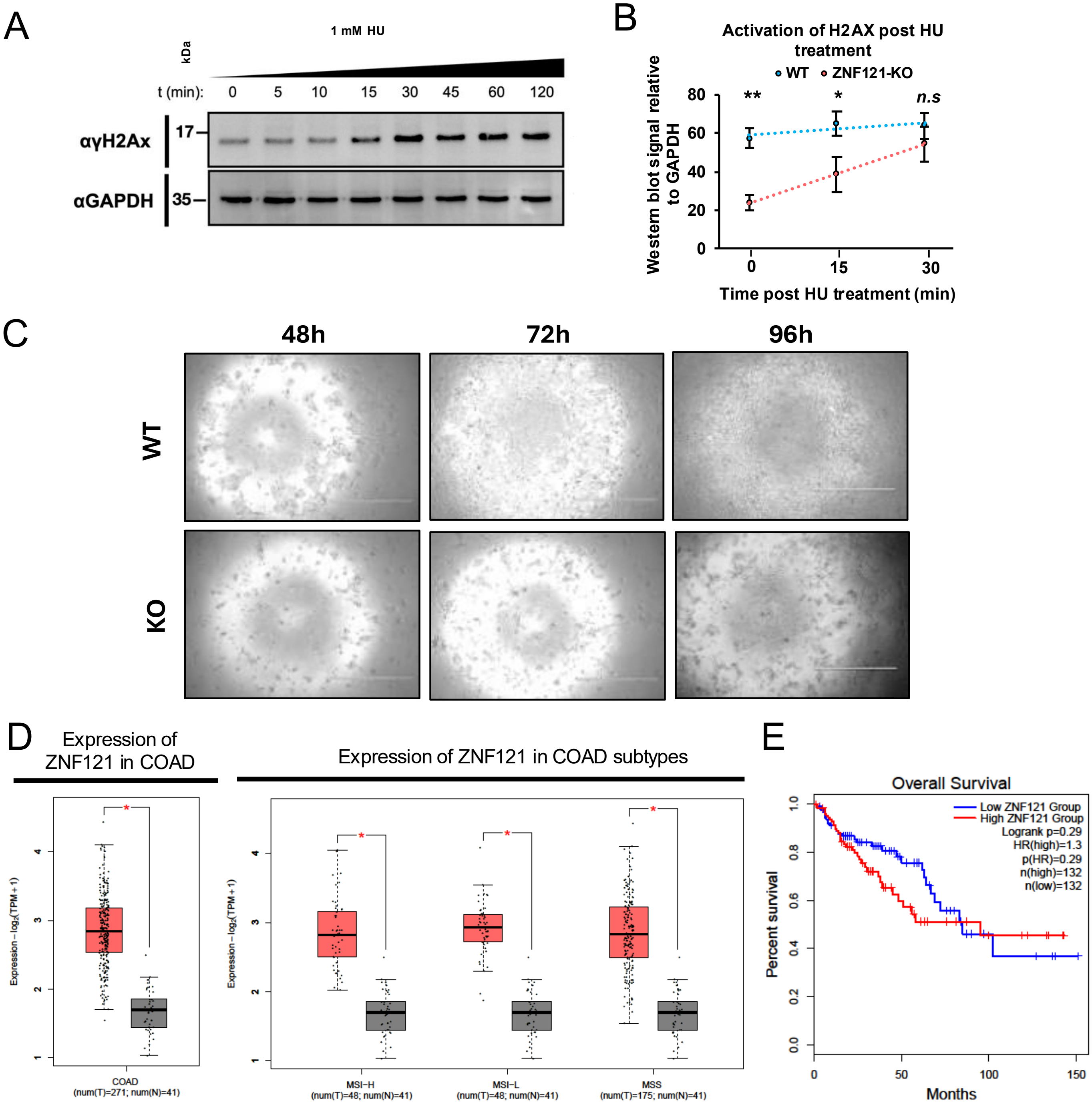
